# Microbiota and metabolic adaptation shape *Staphylococcus aureus* virulence and antimicrobial resistance during intestinal colonization

**DOI:** 10.1101/2024.05.11.593044

**Authors:** Chunyi Zhou, Miranda B. Pawline, Alejandro Pironti, Sabrina M. Morales, Andrew I. Perault, Robert J. Ulrich, Magdalena Podkowik, Alannah Lejeune, Ashley DuMont, François-Xavier Stubbe, Aryeh Korman, Drew R. Jones, Jonas Schluter, Anthony R. Richardson, Paul D. Fey, Karl Drlica, Ken Cadwell, Victor J. Torres, Bo Shopsin

## Abstract

Depletion of microbiota increases susceptibility to gastrointestinal colonization and subsequent infection by opportunistic pathogens such as methicillin-resistant *Staphylococcus aureus* (MRSA). How the absence of gut microbiota impacts the evolution of MRSA is unknown. The present report used germ-free mice to investigate the evolutionary dynamics of MRSA in the absence of gut microbiota. Through genomic analyses and competition assays, we found that MRSA adapts to the microbiota-free gut through sequential genetic mutations and structural changes that enhance fitness. Initially, these adaptations increase carbohydrate transport; subsequently, evolutionary pathways largely diverge to enhance either arginine metabolism or cell wall biosynthesis. Increased fitness in arginine pathway mutants depended on arginine catabolic genes, especially *nos and arcC*, which promote microaerobic respiration and ATP generation, respectively. Thus, arginine adaptation likely improves redox balance and energy production in the oxygen-limited gut environment. Findings were supported by human gut metagenomic analyses, which suggest the influence of arginine metabolism on colonization. Surprisingly, these adaptive genetic changes often reduced MRSA’s antimicrobial resistance and virulence. Furthermore, resistance mutation, typically associated with decreased virulence, also reduced colonization fitness, indicating evolutionary trade-offs among these traits. The presence of normal microbiota inhibited these adaptations, preserving MRSA’s wild-type characteristics that effectively balance virulence, resistance, and colonization fitness. The results highlight the protective role of gut microbiota in preserving a balance of key MRSA traits for long-term ecological success in commensal populations, underscoring the potential consequences on MRSA’s survival and fitness during and after host hospitalization and antimicrobial treatment.

**Importance:** The fitness of MRSA depends on its ability to colonize. A key, underappreciated observation is that gut colonization frequently serves as the site for MRSA infections, especially among vulnerable groups such as children and hospitalized adults. By evolving MRSA strains in germ-free mice, we identify molecular mechanisms underlying how MRSA exploits a depletion in host microbiota to enhance gut colonization fitness. This work points to bacterial colonization factors that may be targetable. Our findings indicate that adaptive changes in MRSA often reduce its antimicrobial resistance and virulence, and are suppressed by the presence of native commensal bacteria. This work helps explain the ecology of pathoadaptive variants that thrive in hospital settings but falter under colonization conditions in healthy hosts. Additionally, it illustrates the potential adverse effects of prolonged, broad-spectrum empirical antimicrobial therapy and adds a new type of weight to calls for microbiota transplantation to reduce colonization by antimicrobial-resistant pathogens.

## INTRODUCTION

Among antimicrobial-resistant bacteria, methicillin-resistant *Staphylococcus aureus* (MRSA) causes the second highest morbidity and mortality worldwide (1). Since colonization is an important prerequisite for *S. aureus* infection and transmission (2), the fitness of MRSA as a pathogen depends on its ability to colonize. A complex relationship exists between colonization, microbial virulence, antimicrobial resistance, and fitness among commensal competitors, thereby complicating the study of the evolution of colonization-adaptive traits. Understanding this relationship has important implications for both pathogen biology and public health.

Nares are the primary site of colonization by *S. aureus* (3), However, gastrointestinal carriage of *S. aureus*, and especially MRSA, is common in some vulnerable host populations. For example, community-acquired (CA)-MRSA frequently colonizes the gastrointestinal tract of infants, and the rectum and perianal skin are key sites for surface colonization in children with CA-MRSA skin infections (4, 5), We recently discovered metabolic changes that promote intestinal colonization in a strain of CA-MRSA primarily afflicting children (6), That pathogen advantage primes the clone for epidemiological success. Gastrointestinal (GI) carriage of *S. aureus* is thought to decrease with host maturity owing to the greater complexity of the microbiota of adults compared to infants, a process known as colonization resistance (7), Nevertheless, gastrointestinal colonization by MRSA is common in hospitalized adults, as disruption of microbiota by antibiotics and critical illness predispose to bacterial overgrowth with *S. aureus* and other pathogens (8–12). One result is fecal shedding that promotes environmental contamination and transmission of MRSA in hospitals. Moreover, rectal carriage of MRSA is associated with an increased rate of invasive infection in high-risk patients (13, 14). Collectively, these and other observations (15, 16), indicate that gastrointestinal colonization in the setting of microbiota dysbiosis contributes to the spread and pathogenesis of MRSA. We currently lack sufficient insight to interrupt this process.

In the absence of growth suppression by gut commensal gut microbes, MRSA overgrowth can occur. To date, little is known about how the bacterium adapts to the intense competition that ensues among MRSA cells. To explore gut adaptation, we examined MRSA colonization of mice. Since microbiota composition is highly variable among mice following antibiotic treatment, patterns of molecular evolution associated with antibiotic treatment during colonization of mice are highly variable (17). Consequently, identification of colonization-adaptive genes is more straightforward in germ-free mice, which provide a simplified, ecologically relevant model to study the molecular basis and fitness effects of adaptive mutations in gut lacking commensal competitors. In this connection, the current study used genomic analyses and competition assays to characterize how the absence of gut microbiota in germ-free mice shape the evolution of MRSA.

## RESULTS

### MRSA rapidly evolves in germ-free mice

To study the evolutionary dynamics of MRSA during adaptation in the GI tract of germ-free mice, we introduced community-acquired MRSA (CA-MRSA) strain USA300 LAC (18) by gavage (inoculum size ∼5 x 10^8^ colony-forming units [cfu]) into four sets of germ-free mice that were housed in independent cages. Stool pellets were harvested at weekly intervals for determination of *S. aureus* cfu (Fig. 1A). Recovery of *S*. *aureus* consistently averaged ∼10^9^ cfu/g stool from week 1 until the end of the experiment (Fig. 1B). For comparison, conventional mice in our facility often remain colonized at much lower levels (∼10^5^ cfu/g stool) (Fig. 1C). Thus, MRSA populations expand to high densities in the intestinal tract of germ-free mice (19).

**Fig. 1.**
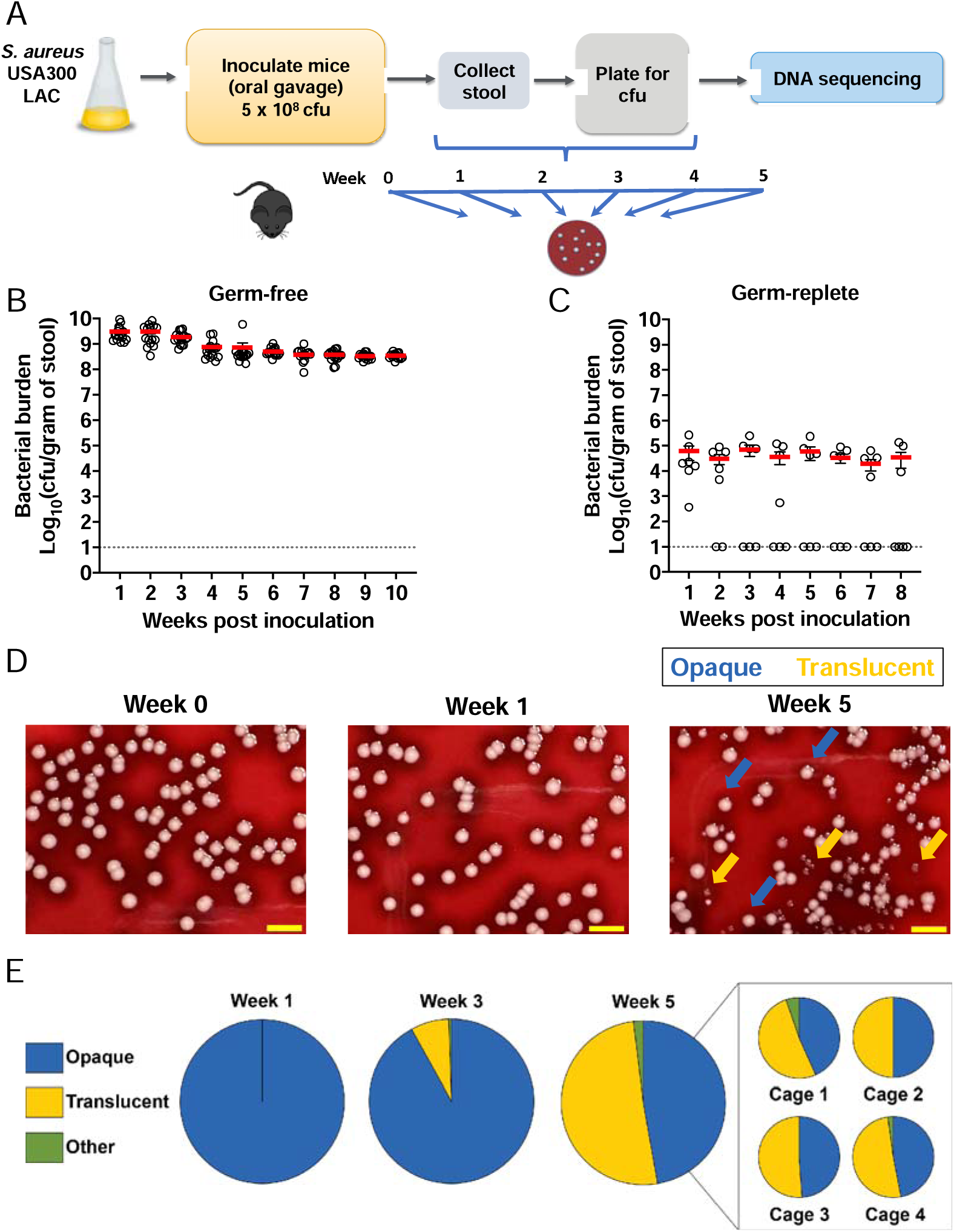
Evolution of morphologic variants in the gut of germ-free mice. (A) Experimental design. Germ-free C57BL/6 mice housed in 4 cages were individually gavaged with 5 × 10^8^ cfu of strain LAC (BS819). (B, C) Quantification of bacteria (cfu) in stool from germ-free (B; *n* = 16) and conventional (germ-replete; C; *n* = 7) mice. Each symbol represents data from one mouse. Data are mean ± SEM. Dotted line, limit of detection. (D) Colony morphology. Representative images at week 0 (LAC wild-type), 1, and 5 (left, middle, and right panels, respectively). Blue arrows, opaque (wild-type-like) colonies; yellow arrows, translucent (variant) colonies. (E) Colony morphology over time. Morphotypes were scored by plating bacteria (*n* > 200 colonies) on tryptic soy agar with sheep blood from all 4 cages at each timepoint. Also shown are week 5 data from individual cages.

After ∼3 weeks, translucent colony variants appeared that were distinct from the opaque morphotype of the wild-type parental strain of the initial inoculum (Fig. 1D-1E). Translucent variants evolved in all four cages in similar proportions, suggesting that their selection was driven by adaptive genetic variations. Translucent variants reached a peak prevalence of ∼50% at five weeks (Fig. 1E), and then their prevalence plateaued, suggesting adaptive changes were also present in the wild-type-like (opaque) phenotype. Since rates of fitness improvement are expected to show diminishing returns with time (20), subsequent analysis focused on week-5 mice to capture most of the gain that improves colonization.

### Adaptation targets a limited set of transcriptional and regulatory elements

To identify the genetic basis of adaptation and link genotype to phenotype, we determined the genome sequence of the parental strain used as inoculum and ∼25 translucent and ∼25 opaque colonies in mice in each of the four cages five weeks post-colonization (*n* = 202 strains; Dataset S1). The majority (74/691 or ∼83%) of mutations identified were point mutations. Among these, most (321/574 or ∼56%) were missense, 144 (∼25%) were synonymous, 14 (∼2%) were translation-stop mutations, and 95 (∼17%) were intergenic. The mean number of mutations per evolved MRSA clone was 3.4, with a minimum of zero and a maximum of ten. The total number of mutations accumulated was between 40 and 54 distinct mutations per cage (including indels and complex mutations). We found no evidence of hypermutation, as evidenced by a relative absence of 1) low-frequency mutations (Dataset S1), 2) an imbalance in favor of transitions over point mutations, and 3) mutations in known mutator genes (e.g., *uvrABC*, *mutS*) (21). Moreover, mutation frequencies, measured by mutation to rifampin resistance, were identical when colonization-adapted (evolved) and parental strains were compared (Fig. S1). Thus, bacterial load and spontaneous mutation explain adaptation.

We identified multiple parallel mutations, defined as any gene, intragenic region, or metabolic pathway carrying mutations in populations from all cages of mice and with a sum of mutation frequency in each mutational target of ≥ 30% in each cage (e.g., sugar transport, arginine metabolism). Targets that fit our criteria included carbohydrate transport *(glcA*/*B*/*T, fruB)*, serine deamination *(sdaA)*, arginine metabolism (*ahrC*/*arcR*), and cell wall metabolism (*walKR)* (Fig. 2A). Colony translucency correlated almost exclusively with *walKR* mutations, indicating a genetic basis for the phenotype. Opaque-colony phenotypes almost always carried mutations in *ahrC*/*arcR*. Additionally, we observed frequent excision of the SCC*mec* element, which contains *mecA* and forms a composite genomic island with the arginine catabolic mobile element (ACME) (22). They encode methicillin resistance and an accessory copy of the arginine deiminase operon, respectively. In three of four cages, the SCC*mec* excision correlated with the opaque phenotype and chromosomal mutations in *ahrC or arcR*, thereby linking changes in chromosomal and accessory arginine metabolism.

**Fig. 2.**
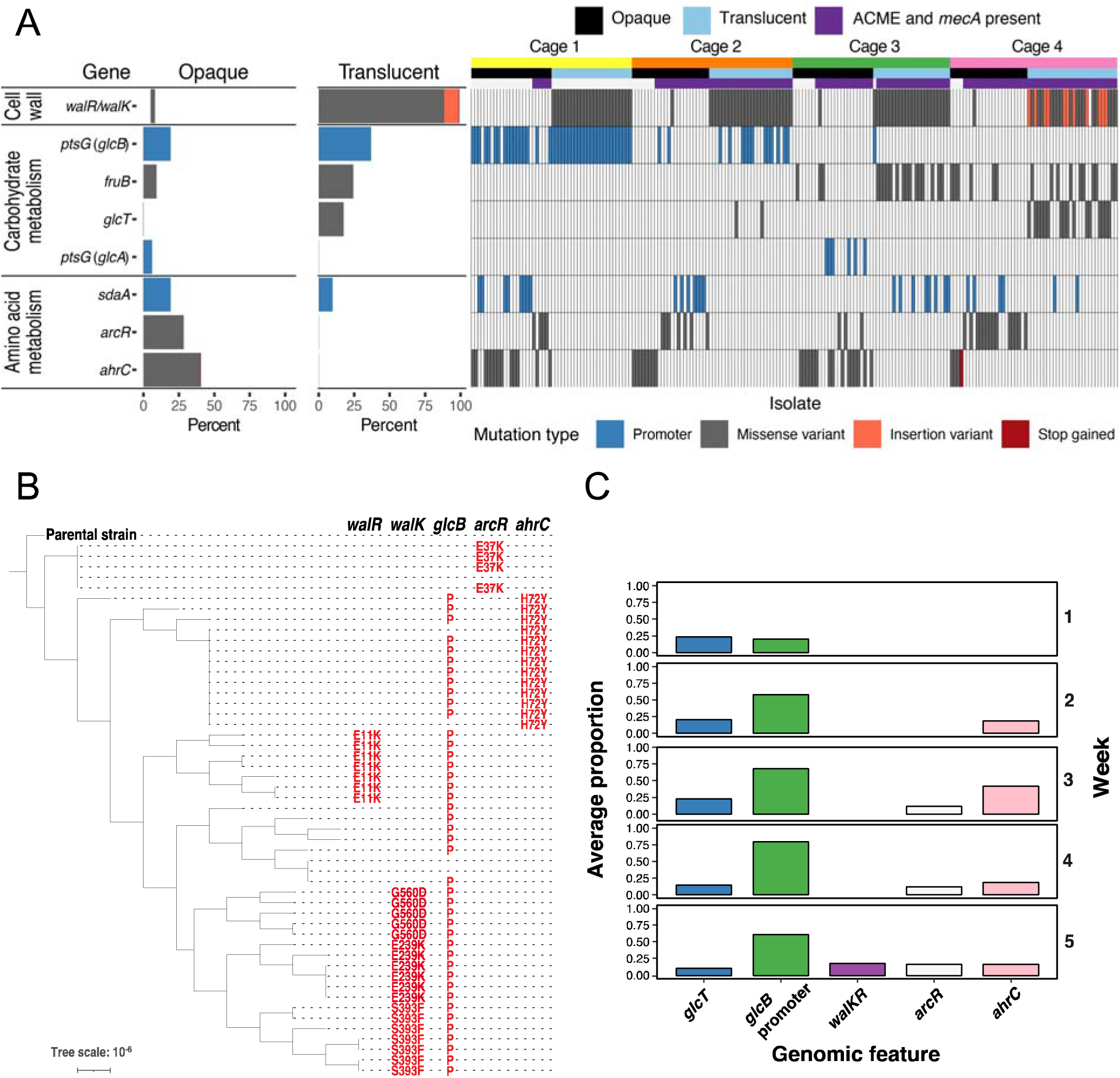
Parallel evolution of mutations and genetic alterations within and between cages of germ-free mice. (A) Distribution of top mutated genes in evolved strains (see Results). Single evolved colonies (*n* = 201) of strain LAC (BS819) from four cages of germ-free mice were sequenced 5-weeks post-inoculation. Mutation type, opaque and translucent morphology, and presence or absence of methicillin-resistance (*mecA*) and ACME (arginine catabolic mobile element) are indicated. Insertion variants are in-frame with the *walK* coding sequence. See Table S1 and Fig. S2 for supporting information. (B-C) Primary differentiation by *ptsG*/*glcT* is for the most part followed by polymorphism of *walKR* or *arcR*/*ahrC*. (B) Phylogeny of evolved mutants. Maximum-likelihood trees of 51 opaque (*walKR* mutant) and translucent (all other) colonies obtained 5-weeks post-inoculation from cage 1 mice. Gene names are listed at the top, and mutations for each isolate are indicated on the corresponding horizontal line and column. (C) Changes in mutation composition over time. Aggregate allele frequency estimates (fraction of aligned reads) in selected genes by week, identified by deep sequencing of thousands of pooled colonies obtained from the stool of mice in all four cages.

A parallel evolution study of strain JH1, which belongs to a highly prevalent hospital-associated (HA-) MRSA lineage (23), revealed a similar mutation pattern, including repeated selection of *walKR* and *glcB* mutations, although differences in mutations that control enzymes involved in arginine metabolism were observed (e.g., enrichment of mutations in *rsaE*, a regulatory RNA that controls enzymes involved in arginine metabolism (24), and a concurrent decrease in *ahrC/arcR* mutations)(Dataset S2). Thus, dominant adaptations were not MRSA strain-specific.

### Sequential evolution of adaptive mutations

Phylogeny of the evolved LAC variants revealed a deeply branched tree with multiple lineages (Fig. 2B). Lineage-specific diversity corresponded largely to mutually exclusive mutations in *ahrC, arcR,* or *walKR*. In contrast, identical mutations in *glcB/glcT* generally occurred throughout the phylogeny. Collectively, these observations suggest that, for the most part, primary differentiation of *glcB/glcT* is followed by polymorphism of either *ahrC/arcR* or *walKR*.

To determine the time of the emergence of these mutations, we deep sequenced populations of strain LAC by pooling clones sampled from each cage weekly for five weeks (Fig. 2C). Deep sequencing involved pooling hundreds of colonies from each sample and sequencing the pool with high coverage (∼450x). Glucose transport mutations in *glcB* were detected in the first week post-colonization and thereafter (Fig. 2C and Dataset S3). Mutations in *ahrC* and *arcR* were initially detected in the 2^nd^ and 3^rd^ week, respectively; *walKR* mutants appeared in the 5^th^ week (Fig. 2C). In contrast, analysis of single colonies discussed above identified translucent *walKR* mutants in the 3^rd^ week of evolution (Fig. 1E), likely owing to the enhanced sensitivity of analyzing individual colonies for identifying mutations having lower allele frequencies.

Nonetheless, the combination of deep and single colony sequencing results supports the idea that LAC adapts rapidly to the intestine of germ-free mice by mutating glycolytic pathways, followed by repeated selection of mutations that reprogram arginine or cell wall metabolism.

### Adaptation is microbiota-dependent

To confirm that evolved changes in strain LAC are adaptive, we competed a 1:1 mixture of a cadmium resistance-marked parental strain with evolved mutants by colonizing germ-free mice. The following evolved mutants were evaluated: 1) a *glcB* mutant that lacked mutations elsewhere in the genome, 2) an *ahrC*/ACME double mutant, and 3) a *glcB*/*walK* double mutant. All of the evolved strains, especially the *ahrC*/ACME double mutant (100-1000 fold) and *glcB*/*walK* double mutant (10-100 fold), outcompeted wild-type (Fig. 3A, 3C, and 3E). The fitness of the marked parental strain *in vivo* matched the unmarked parental strain; thus, results were not skewed by an effect of the resistance marker (Fig. S3A-S3B).

**Fig. 3.**
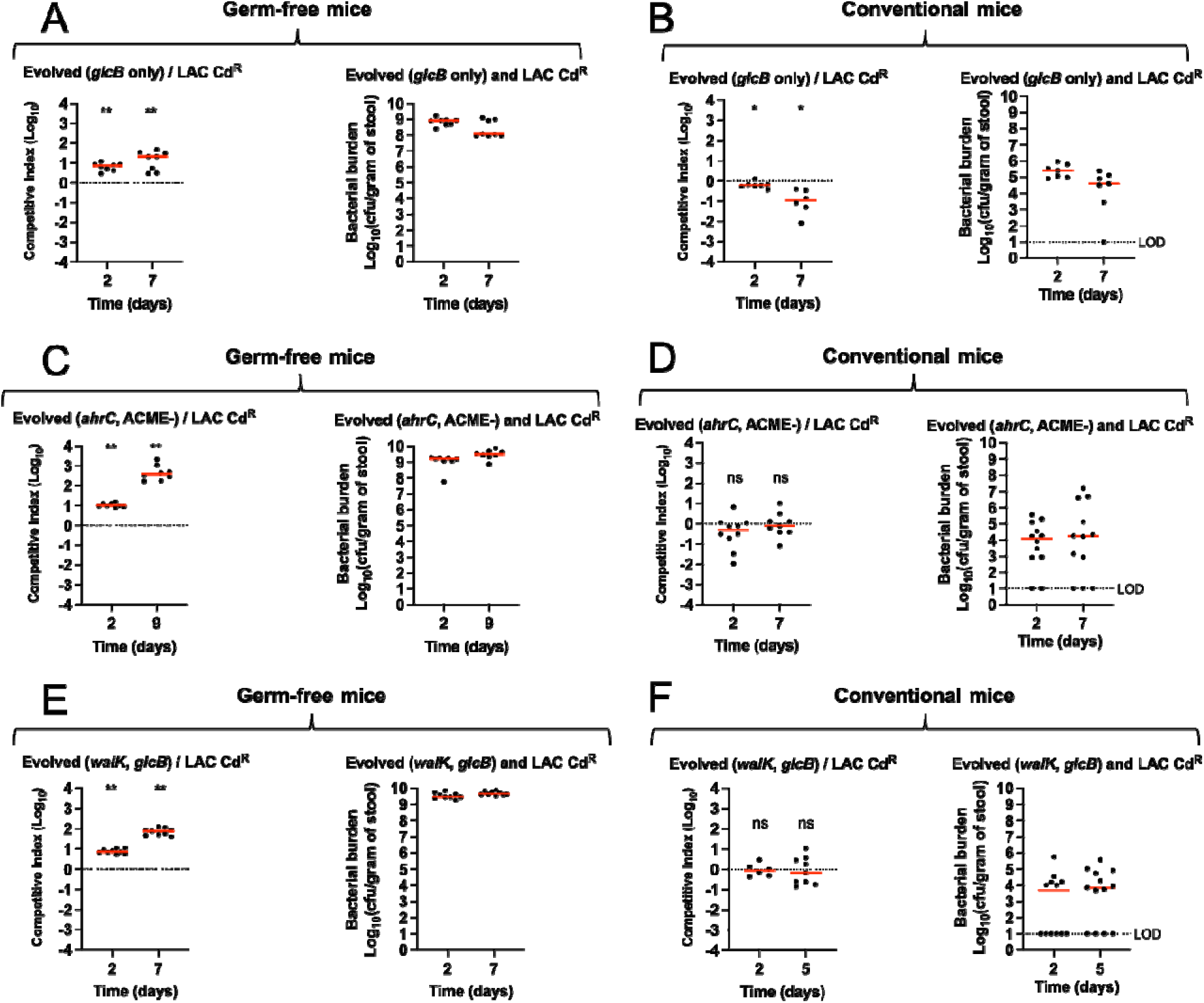
Fitness of evolved mutants in the gut of germ-free and conventional mice. **(A-F)** Competitive colonization assays (competitive index, *Left*) and quantification of bacteria in stool (cfu, *Right*) determined from the stool of colonized germ-free (A, C, and E) or conventional, germ-replete mice (B, D, and F). Strains with evolved mutations in *glcB* (BS1565; panels A and B), *ahrC* ACME (-) (W5-1-O18; panels C and D), and *walk glcB* (W5-9; panels E and F) were competed against patental strain LAC. LAC contained a chromosomally integrated cadmium resistance marker (SaPI1 *att*C::*cadCA;* strain VJT32.58) to distinguish the strains following plating of serial dilutions on tryptic soy agar (TSA) with or without cadmium (0.3 mM). Evolved mutants and LAC Cd^R^ were mixed 1:1 and used to inoculate each mouse. Each symbol represents data from one mouse (*n* = 6-12 mice). Median values (red lines) are shown, and each symbol is the competitive index (*Left*) or cfu (*Right*) from one mouse. \**P* < 0.05, \*\**P* < 0.01, and ns (*P* > 0.05) by Wilcoxon signed-rank tests. The dotted lines indicate a 1:1 ratio (equal fitness). LOD: limit of detection.

Notably, we found that evolved mutants did not display significantly increased GI competitive fitness compared to WT in germ-replete (conventional) mice (Fig. 3B, 3D, and 3F). Moreover, genome sequencing of randomly selected week-5 colonies (*n* = 10 colonies) in conventional mice showed that MRSA does not acquire mutations during colonization of conventional mice. To determine whether a ‘low-complexity’ microbiota, which might be relevant to antibiotic-induced disruption of microbiota, is more permissive for adaptation, germ-free mice were colonized with minimal defined flora consisting of a consortium of 15 bacterial strains representing the murine gut microbiota(25) followed by strain LAC 3 days later. The consortium is known to confer resistance to colonization by microbial pathogens in mice (25). As with conventional mice, we found no mutation after colonization for 5 weeks (10 colonies sequenced). Collectively, the results indicate that commensal microbiota limits both strain LAC colonization and adaptation.

### Adaptation supports MRSA growth and biomass

#### Increased glycolytic activity

Mutation of the *glcB* (*ptsG*; SAUSA300_2476) promoter was among the most common recurring events, being present in 28.4% (57 of 201) of all evolved clones of strain LAC at 5 weeks (Dataset S1, Fig. 2, and Figs. S2A and S4A). Unlike the 3 other phosphoenolpyruvate-dependent phosphotransferase system (PTS) glucose transporters in *S. aureus* (26), expression of *glcB* is thought to be induced rather than constitutive, potentially explaining why its regulatory region mutated preferentially. The upstream regulatory region of *glcB* contains a putative terminator structure overlapping a conserved ribonucleic anti-terminator sequence (RAT) motif (Fig. S4B-S4C). In *Bacillus subtilis* and *Staphylococcus carnosus* (27, 28), and likely *S. aureus,* the RAT motif is thought to be recognized by the transcriptional anti-terminator protein GlcT, which was another target of adaptive mutation. In the absence of glucose, a conserved histidine residue (His-104) in GlcT is phosphorylated, thereby inactivating the protein. If glucose is present, unphosphorylated GlcT binds to RAT, which prevents formation of the terminator structure, thereby enabling transcription of the downstream transporter gene (Fig. S4B). 80% (45 of 57) of clones containing mutations in *glcB* had mutational hotspot deletions in RAT that are predicted to inhibit formation of the terminator structure (Fig. S4B-S4C). Additionally, mutations in *glcT* were in all cases confined to a hotspot in the phosphorylation site (His-104), which would constitutively activate GlcT (Fig. S4B). Thus, both mutations are expected to de-repress *glcB* and/or other glucose transporters. Indeed, *glcB* expression in independently evolved strains containing either *glcB* or *glcT* mutations was low in the presence of glucose (14 mM) but was ∼300-fold higher compared to the parental strain when glucose was absent (Fig. 4A-4B). We conclude that the evolved mutants constitutively express *glcB* and that His-104 in GlcT is likely required for negative regulation of the antiterminator activity.

**Fig. 4.**
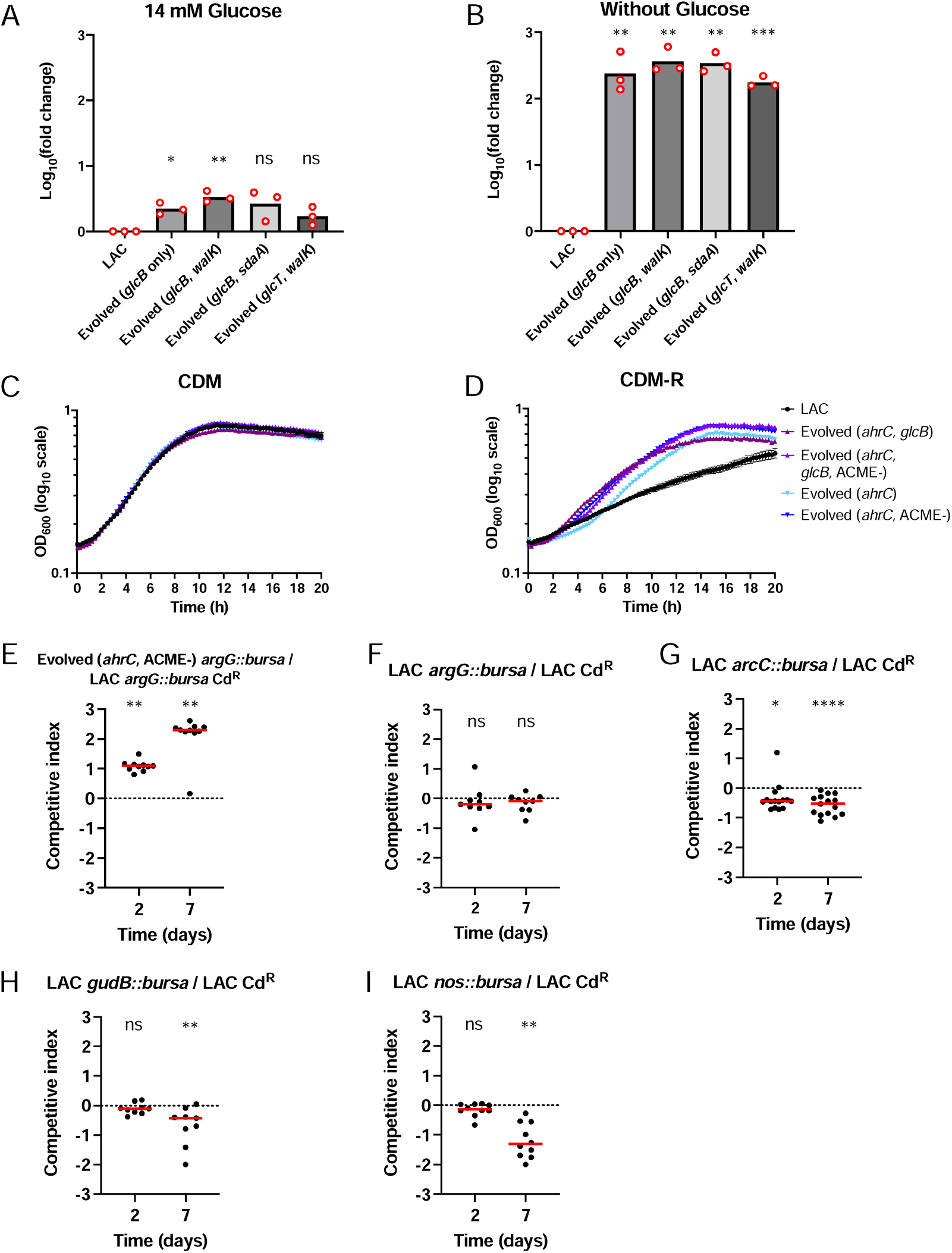
Involvement of evolved mutations with upregulation of glucose import and arginine catabolism. (A-B) Effect of evolved mutations on transcription of *glcB* in the presence and absence of glucose. Total cellular RNA was extracted from the indicated evolved mutant or parental strain LAC (BS819) after aerobic growth in chemically-defined medium (CDM, 1% cas amino acids with 14 mM glucose) (A) or CDM without glucose (B), followed by reverse transcription and PCR amplification of *glcB*, using 16S rRNA as an internal standard. Data are mean log_10_(fold change) from each strain (*n* = 3). ns *P* > 0.05; * *P* < 0.05; ** *P* < 0.01; *** *P* < 0.001 by one sample *t* test comparing to 0 (log10 [LAC fold change]). Strains were evolved mutant *glcB* (BS1565), *glcB walK* (W5-9), *glcB sdaA* (W5-10); and *glcT walK* (W5-17). (C-D) Effect of evolved mutations in *ahrC* on growth in media lacking arginine. Growth analysis of evolved mutants *ahrC glcB* (W5-7), *ahrC glcB* ACME (-) (W5-1-O1), *ahrC* (W5-2-O10), *ahrC* ACME (-)(W5-1-O18), and parental strain LAC (BS819) in CDM with or without (CDM-R) arginine. Data represent means ± SEM from three (*n* = 3) biological replicates. (E-I) Effect of transposon insertions in arginine biosynthesis and catabolism genes in evolved strains. (E-I) Competition assays in germ-free mice, performed as in Fig. 3, involving (E) evolved mutant *ahrC* ACME (-) (W5-1-O18) containing arginine biosynthesis gene mutation *argG::bursa* (BS1541) or LAC *argG::bursa* Cd^R^ (BS1539), and (F-I) parental strain LAC Cd^R^ (VJT32.58) and arginine catabolic pathway transposon mutants *arcC::bursa* (BS1535), *gudB::bursa* (BS1409), and *nos::bursa* (BS1434). See Fig. S3 for bacterial burden. Each symbol represents data from one mouse (n = 9-15 mice). Wilcoxon signed-rank test: ns *P* > 0.05; * *P* < 0.05; ** *P* < 0.01; **** *P* < 0.0001. The red lines are medians.

As with *glcB*, mutations in *sdaA* consisted almost exclusively of hotspot mutations in the upstream promoter region (Fig. S2A). *sdaA* catabolizes L-serine to pyruvate, the final product of glycolysis, to synthesize ATP via substrate-level phosphorylation (29, 30). The ability to catabolize L-serine increases bacterial fitness by providing *Enterobacteriaceae* with a growth advantage in inflamed gut (31). The mutations in *sdaA* are likely upregulating since attenuating mutations would be far more frequent in coding regions. This finding, together with the above mentioned increased *in vivo* fitness of evolved *glcB* and *glcB sdaA* double mutants (Fig. 3A), supports the idea that upregulating mutations in glycolytic pathways are selected during GI adaptation in the absence of gut microbiota.

In 3 cages of mice, mutations in *glcB/T* and/or *sdaA* were mutually exclusive with mutations in *fruB* (fructose 1-phosphate kinase); in one cage *fruB* mutations were the dominant mode of carbohydrate metabolism mutation (Fig. 2A, Dataset S1). Thus, glucose transport is not the only carbohydrate metabolic pathway targeted by mutation during adaptation to the GI tract (Fig. S2B).

#### Increased arginine catabolism

Following mutation of glycolytic pathways, opaque phenotypes primarily accumulated mutations in arginine metabolic genes *ahrC* or *arcR*. *ahrC* represses the arginine biosynthetic pathway and the arginine deiminase (ADI) operon; *arcR* upregulates the ADI pathway (32, 33)(Fig. S2B). Mutations in these two pathways were mutually exclusive, suggesting that mutation of either is functionally redundant and presumably the result of convergent evolution. Recent work demonstrated that mutations in *ahrC* that facilitate arginine biosynthesis are selected in clinical infections (33). We therefore focused our analysis on *ahrC* mutants.

*S. aureus* is an arginine auxotroph, but inactivation of *ahrC* upregulates *argGH* (argininosuccinate synthase/lyase) and *arcB1* (ornithine carbamoyltransferase), thus enabling arginine biosynthesis via proline in media lacking glucose (glucose suppresses arginine synthesis) (33). Gut-evolved *ahrC* mutants grew robustly in media lacking arginine and glucose, phenocopying an engineered *ahrC* deletion mutant (33) and indicating that the mutations are inactivating (Fig. 4C-4D). However, mutation of the arginine biosynthesis pathway (*argG::Tn)* in the evolved *ahrC* mutant and parental backgrounds failed to eliminate the *ahrC* competitive advantage *in vivo* (Fig. 4E), indicating that *ahrC*-mediated fitness in the GI tract of germ-free mice does not require arginine biosynthesis. Consistent with this idea, the inactivation of *argG* in the wild-type background had no effect on competitive fitness (Fig. 4F). Thus, *ahrC* must exert its fitness-enhancing effect through arginine catabolism.

*S. aureus* employs three pathways to catabolize arginine: 1) the arginine deiminase pathway, which generates substrate for ATP via carbamoyl phosphate (34), 2) the arginase pathway that produces glutamate, which replenishes the tricarboxylic acid cycle via 2-oxoglutarate, and 3) nitric oxide synthase (*nos*), which converts arginine to nitric oxide and ultimately nitrite to nitrate for use in respiration and adaptation to low-oxygen environments (35). To determine which catabolic pathway is important for colonization *in vivo*, we performed *in vivo* competition experiments using mutants from each pathway (*arcC*, carbamate kinase; *gudB*, glutamate dehydrogenase; *nos*, nitric oxide synthase) from a sequence-defined transposon library (36) (Fig. 4G-4I). All 3 mutants were outcompeted by wild-type, but *nos* inactivation had by far the most profound (∼20 fold on day 7) detrimental effect. Collectively, these data support the ideas that 1) arginine catabolism is critical for enteric fitness in germ-free gut, and 2) mutation of *ahrC* functions to increase arginine catabolism.

The role of adaptive mutations in other arginine metabolism genes, such as *arcR*, ACME, spoVG, and *rsaE* (in strain JH1), requires additional study. However, mutations in *spoVG* and the ACME-encoded *ahrC* homolog *argR2*, like those in native *ahrC*, facilitate growth in media lacking arginine and glucose (33). Relatedly, excision of SCC*mec*, which was a frequently observed adaptation in strain LAC (31%) (Fig. 2A), occurred only three times in JH1 (3%). SCC*mec* in JH1 does not encode ACME as it does in LAC (Dataset S2). These observations suggest that selection against ACME-mediated effects on arginine metabolism drives SCC*mec* excision in strain LAC.

### MRSA shapes fecal metabolite concentrations

To determine whether MRSA causes a shift in the fecal metabolome that correlates with evolved mutations, we collected stool pellets from our evolution experiment a week after colonization for metabolite profiling. Metabolomic profiling showed a distinct clustering pattern between pre-inoculated germ-free mice (week 0) and those given LAC (week 1) (Fig. 5A).

**Fig. 5.**
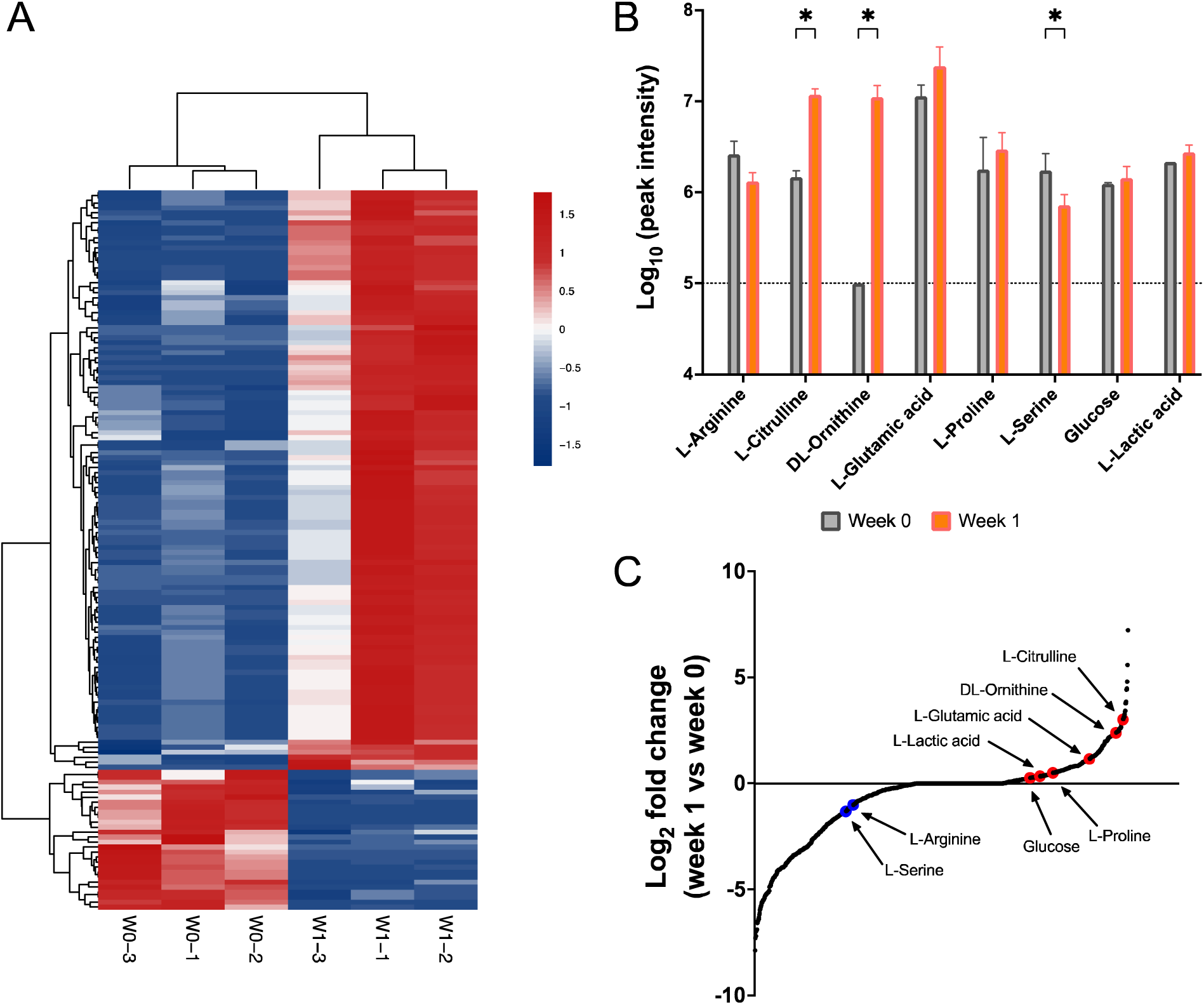
MRSA shapes fecal metabolite concentrations. Comparative analysis of the fecal metabolome of germ-free mice (*n*lJ=lJ3) housed in a single cage before (week 0; W0) and after (week 1; W1) inoculation with parental strain LAC (BS819). The homogenized stool (total 3 stool pellets per week) was analyzed by HILIC UPLC mass spectrometry. (A) Metabolite profile comparison of week 0 and week 1 stool. Unsupervised clustering analysis showing significant altered metabolites (*P* < 0.05 from *t* test; *n* = 144). (B) Relative concentrations of metabolites associated with carbohydrate and arginine metabolism. Peak spectra intensities of the indicated metabolites from week 0 and week 1 stool samples. Data are mean ± SD. The dotted line indicates the limit of detection. Unpaired *t* test, * *P* < 0.1. (C) Log_2_ fold change of all metabolites (*n* = 1320) in stool samples from week 1 relative to week 0. Each dot represents a metabolite. Red and blue indicates metabolites that were increased or decreased, respectively.

Glucose and arginine are of particular interest, because their metabolic pathways were targeted by mutation during gut colonization (Fig. S2A-S2B). We found that metabolites identified as the arginine breakdown products ornithine and citrulline were increased when LAC was present (Fig. 5B-5C). These data, together with competition assays providing direct evidence that evolved mutants utilize arginine more efficiently for growth than parental strain LAC (Fig. 4), suggest that increases in metabolites represent MRSA-derived products of arginine catabolism. The observed increase in arginine metabolites was accompanied by a small decrease in arginine levels (Fig. 5B-5C). Serine levels also decreased when LAC was present (Fig. 5B-5C), consistent with conversion of serine to pyruvate for energy via promoter mutation of *sdaA* (Fig. S2A).

Examination of the fecal metabolome in subsequent weeks indicated consistent increases in arginine and glucose catabolic products, stability of arginine levels, and a slight decrease in glucose. GI tract concentrations of arginine and other amino acids are known to be elevated in germ-free mice (37, 38). Thus, the relative absence of arginine or glucose depletion supports the idea that evolved MRSA outcompete the parental wild-type by competition for glucose and arginine that are abundant because there are no commensal bacteria in the intestine to consume them.

### *walKR* mutations increase biomass and enteric fitness

Evolved mutations also occurred in genes frequently mutated in vancomycin intermediate-resistant *S. aureus* (VISA) that belong to the essential two-component system *walK* and *walR*. This system links cell wall biosynthesis to cell division (39). *walKR* mutations were tightly linked to members of the translucent colony class (Fig. 2A), indicating that they are the basis of the phenotype.

Only missense and insertion mutations in *walKR* were observed; frameshift or nonsense mutations were absent (Fig. 2A and Fig. S2A). Additionally, mutations did not match any known VISA-associated mutation (40), which usually attenuate *walKR* activity. Thus, mutation patterns suggest that evolved mutations upregulate activity of the operon. Consistent with this idea, evolved mutants demonstrated increased autolysis and biofilm biomass (Figs. 6A and B), which are positively controlled by *walKR* (39). However, the test strain, like all *walKR* mutants, contained mutations in other genes, primarily *glcB*. To rule out the effects of pleiotropy from other mutations in the strain background, we constructed a site-specific replacement (H271Y) in WalK of the parental strain LAC. This mutation is known to activate the *walKR* regulon (41). As predicted, the substitution increased vancomycin susceptibility, autolysis, and colonization fitness (Fig. 6) compared to wild-type LAC. These findings support a direct relationship between colonization fitness, the evolved mutations, and *walKR* activity.

**Fig. 6.**
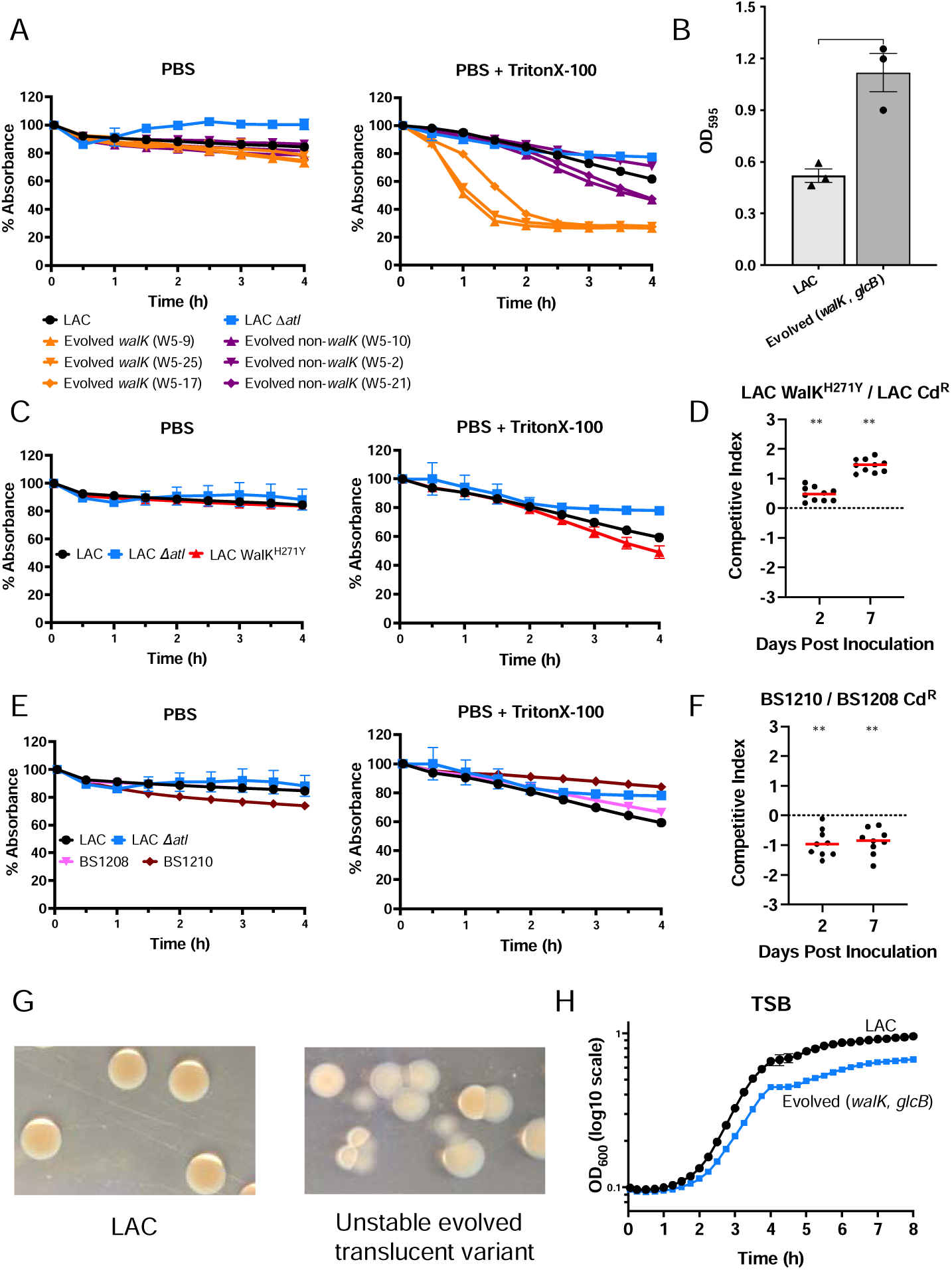
Association of evolved *walKR* mutations with increased autolysis, biofilm, intestinal fitness, and colony morphology. (A) TritonX-100-stimulated autolysis of *walKR* mutant cells. Cells of the indicated evolved *walKR* mutant, parental strain LAC (BS819), and control strain LAC *Δatl* (VJT80.33) grown in tryptic soy broth (TSB), were suspended in PBS or PBS containing 0.1% TritonX-100. Rate of autolysis was monitored as a decrease of absorbance at 600 nm. Data represent the means ± SEM from (*n* = 2-3) biological replicates. (B) Biofilm formation. Biofilm formation in TSB supplemented with 0.25% w/v D-(+)-glucose medium at 37°C for evolved strain *walK*, *glcB* (W5-9) and parental strain LAC with tissue culture-treated 96-well plates. Data represent the mean ± SEM from (*n* = 3) biological replicates. Unpaired t test: ** *P* < 0.01. (C) TritonX-100-stimulated autolysis of LAC WalK^H271Y^ cells. Cells of LAC WalK^H271Y^, LAC, and LAC *Δatl* were analysed as in A. (D) Competition assays in germ-free mice, performed as in Fig. 3, involving LAC WalK^H271Y^ and LAC Cd^R^ (*n* = 10 mice). ** *P* < 0.01 by Wilcoxon signed-rank test. The red lines are medians. See Fig. S3 for bacterial burden. (E) TritonX-100-stimulated autolysis of a *yycH* mutant clinical isolate with a VISA phenotype. Cells of ancestral clinical isolate (BS1208), evolved *yycH* mutant (BS1210), LAC, and LAC *Δatl* were analysed as in A. BS1208 and BS1210 are isolates JH1 and JH6, respectively, in Mwangi et al.^25^ (F) Competition assays in germ-free mice, performed as in Fig. 3, involving BS1210 and BS1208 Cd^R^ (SaPI1 *att*C::*cadCA;* strain BS1709)(*n* = 9 mice). ** *P* < 0.01 by Wilcoxon signed-rank test. The red lines are medians. See Fig. S3 for bacterial burden. (G) Intracolonial phenotypic variation in a *walKR* mutant seen as colony sectoring. Photographs of colonies, illuminated by oblique and transmitted light, derived from control strain LAC and a translucent *walK* variant (W3-1C) grown on TSA. (H) Growth curves. Evolved *walK glcB* mutant (W5-9) and parental strain LAC cultures were grown in TSB following 1,000-fold dilution of overnight cultures. Growth of diluted cultures was monitored for 8 hours every 15 min by measuring the OD_600_ using a Bioscreen C.

Although the majority of translucent colony variants remained stable when grown *in vitro*, indicating a heritable change, a small number reverted with passage or demonstrated sectored colonies (portions of the colony reverted to an opaque morphology) (Fig. 6G). The presence of revertants indicates a loss of growth fitness *in vitro*, suggesting tradeoffs in bacterial fitness.

Indeed, *walKR* mutants grew poorly in planktonic cultures (Fig. 6H). Thus, the fitness of evolved *walKR* mutants compared to wild-type *in vivo* was not attributable to an intrinsic growth advantage; if anything, they showed a growth defect.

Mutations that negatively affect the activity of the *walKR* locus are associated with vancomycin-intermediate resistant (VISA) phenotypes (39, 42, 43) and low virulence (39, 44–46). To evaluate colonization phenotypes in such strains, we assayed a clinical heterogeneous VISA (hVISA) clone and its vancomycin-susceptible (VSSA) parent isolated from the bloodstream of a patient with endocarditis who was treated extensively with vancomycin.

Genomic sequencing showed that the variant arose from a common recent ancestor and traced the likely basis of resistance to an attenuating mutation in the *walKR* regulator *yycH*, a frequent target of *walKR-*attenuating mutations in patients (42). As expected, the VISA/*yycH* mutant had decreased autolysis compared to the parental VSSA isolate (Dataset S4). Moreover, when the VISA and VSSA isolates competed in germ-free mice, the resistant variant displayed a colonization defect (Fig. 6E-6F). Thus, *walKR* mutations that decrease susceptibility to vancomycin are attenuated during colonization.

### Colonization adaptation correlated inversely with antimicrobial resistance and virulence

As mentioned above, mutations that decrease *walKR* activity are associated with VISA phenotypes (39, 42, 43). Thus, we were not surprised to find that vancomycin prevents growth of several evolved mutants having *walK* upregulation mutations at drug concentrations that are subinhibitory for wild-type cells (Fig. 7A). At the same time, population analysis for the *walK* mutants, in which large numbers of cells from a culture were applied to antibiotic-containing agar plates and resistant colonies were counted, indicated a similar frequency of colonies that reflect the vancomycin-resistant mutant subpopulations present in the culture (Fig. 7B). Thus, evolved *walK* mutations increased drug susceptibility without interfering with the stepwise accumulation of additional mutations.

**Fig. 7.**
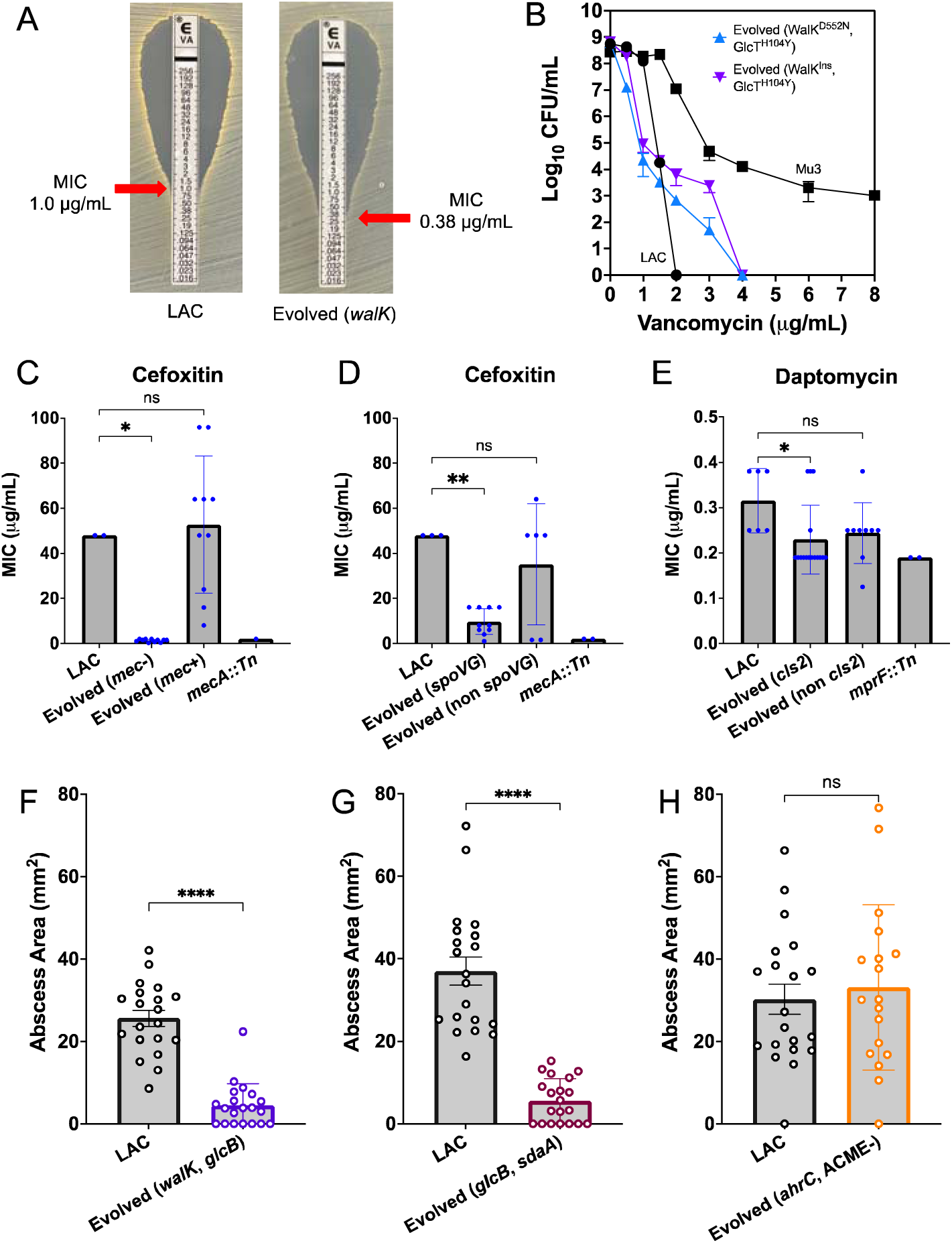
Association of colonization adaptative mutations with antimicrobial resistance and virulence. (A) Vancomycin resistance determined by Etest or population analysis. Minimal inhibitory concentration (MIC) of parental strain LAC (BS819) and an evolved *walK* mutant (W5-2-T2) by Etest. (B) population analysis of LAC, evolved WalK^D552N^ GlcT^H104Y^ mutant (W5-2-T6), evolved WalK^Insertion^ GlcT^H104Y^ mutant (W5-4-T16), and heteresistant control strain Mu3 (BS626). Data represent means from four technical replicates. (C-E) Cefoxitin and daptomycin MICs of parental strain LAC *mecA::bursa* (BS1168), LAC *mprF::bursa* (BS1328), and evolved strains with *mecA*, *spoVG,* or *cls2* mutations, determined by Etest (*n* = 6-16 for evolved strains in each condition). See Table S5 for stain information. ns *P* > 0.05; * *P* < 0.05; ** *P* < 0.01 by one-way ANOVA, Dunnett’s multiple comparisons test. Data are mean ± SEM. (F-H) Association of evolved mutations and skin abscess size. Abscess size, measured 48h after s.c. infection with ∼1 × 10^7^ cfu of the indicated strain, (*n* = 9-10 mice per group with two abscesses per mouse). Strains were evolved mutants *walK glcB* (W5-9), *glcB sdaA* (W5-10), and *ahrC* ACME (-) (W5-1-O18), and parental strain LAC (BS819). ns *P* > 0.05; **** *P* < 0.0001 by Mann Whitney test (F and G) or unpaired *t* test (H). Data are mean ± SEM.

Adaptive mutations in LAC or JH1 also included *mecA* deletions 1) by means of the SCC*mec* excision that eliminates methicillin-resistance and 2) by mutations in antimicrobial resistance loci that are hotspots for adaptation to key anti-staphylococcal antibiotics: *spoVG* (*n =* 25 in strain LAC; oxacillin (47)), *cls2* (*n =* 26 in strain LAC; *n* = 8 in strain JH1; daptomycin), *mprF* (*n* = 9 in strain JH1; daptomycin (48)), and *rpoB* (n = 1 in strain LAC; multiple antimicrobials (49)). We confirmed that 1) *walKR* mutants were more susceptible to vancomycin, 2) *mecA*-deletion mutants were oxacillin susceptible, 3) *spoVG* mutations increased susceptibility to beta-lactams, and 4) *cls2* mutations increased susceptibility to daptomycin (Fig. 7C-7E). Thus, several antibiotic resistance loci are lost or compromised during colonization adaptation.

To determine whether evolved mutations modulate virulence, we compared the virulence of strain LAC to that of evolved *glcB*/*walK*, *glcB*/*sdaA*, and *ahrC/*ACME mutants, which were representative of the dominant evolutionary lineages, in a murine skin abscess model of infection (6). Evolved *glcB*/*sdaA* and *glcB*/*walK* strains formed markedly smaller, or no abscesses, compared with the wild-type parental strain (Fig. 7F-7G). Thus, evolved mutations that enhance colonization attenuate antimicrobial resistance and virulence. In contrast, the parental strain and the evolved *ahrC/*ACME mutant showed similar virulence (Fig. 7H).

Core genome-encoded toxins play an important role in MRSA skin infection (50), and cytotoxicity measurements can be used to determine the potential for MRSA strains to cause disease (51). To determine whether evolved mutations attenuate cytotoxicity, we obtained cell-free extracts from cultures of evolved *glcB*/*walK*, *glcB*/*sdaA*, and *ahrC/*ACME (-) mutants for use in cytotoxicity assays. Evolved clones tended to have equal cytotoxicity toward primary human neutrophils compared to the highly cytotoxic parental strain (Fig. S5). Additionally, exoprotein abundances between the parental strain and the evolved mutants showed, if anything, an increase in exoprotein secretion. Thus, in vitro analyses of cytotoxicity do not correlate with the reduced virulence of colonization-adapted strains.

### Evolved mutations in human gut

To explore the ecological significance of the targets of these mutations within the human gut, we evaluated variant alleles of murine gut adaptive pathways in 395 human gut metagenome samples obtained from an observational cohort of 49 hematopoietic stem cell transplant patients (52). Disruption of gut microbiota in these patients, induced by antibiotics and critical illness, fosters bacterial overgrowth with *Staphylococcus* and other pathogens, facilitating analysis of molecular adaptation (53). Our analysis focused on dominant adaptations that increase glucose transport (*glcB* promoter, *glcT*), arginine biosynthesis (*ahrC*, *arcR*, and *spoVG*), and cell wall metabolism (*walKR*). Additionally, we sought mutations in the chromosomal arginine-deiminase system (Arc) promoter and *ccp*A that, like those in *ahrC* and *spoVG* facilitate arginine biosynthesis (33, 54). Notably, these mutations were recently found to be selected during clinical infection (33).

We detected non-synonymous variant alleles in one sample whose metagenomic assemblies included more than 1 Mbp of *S. aureus* nucleotide sequence, allowing for the assembly of candidate genes (Dataset S5). The sample contained mutations in *spoVG*, one of which was predicted to be deleterious ([e.g., intolerant by the Sorting Intolerant From Tolerant program (55)). This sample also contained a frameshift mutation in *ccpA* and three *arc* promoter mutations. These specific promoter mutations were identical to those found in human infection-associated strains that evolved independently in distinct hosts (33). Moreover, variant *arcA1* promoter mutations co-occurred with variants in other genes (e.g., *spoVG*, *ccpA*) affecting the arginine metabolic pathway, a phenomenon observed during infection (33). This observation suggests genetic instability, potentially facilitating adaptation when selection favors a combination of individually neutral or deleterious mutations.

## DISCUSSION

We report that colonization of germ-free mice by MRSA rapidly selects for the stepwise emergence of bacterial mutations that (*i*) increase carbohydrate transport, thereby priming clones for success; and (*ii*) target two mutually exclusive genetic pathways involving either increased arginine catabolism or cell wall metabolism (Fig. S6A). Notably, the dominant mutants that evolved showed reduced virulence during invasive infection and increased antimicrobial susceptibility. Moreover, we found that microbiota inhibits MRSA metabolic adaptation, thereby maintaining wild-type MRSA characteristics. The interaction of colonization pathways that affect virulence and resistance with fitness among commensal competitors provides a general framework for understanding the ubiquitous presence of wild-type phenotypes in MRSA populations (Fig. S6B).

The scarcity of *S. aureus* sequences in our human metagenomic dataset likely stem from the widespread use of intravenous vancomycin in almost all patients and the empirical administration of oral vancomycin to eradicate *Clostridioides difficile* in approximately one quarter of patients (52). Thus, a broader assessment of additional patients is needed to understand the prevalence and evolution of *S. aureus* variants in the gut. Despite limited samples, observed arginine metabolic mutations imply that *S. aureus* variants in the human gut may alter pathways in a manner similar to those observed in mice. This finding, combined with previous work by others (33), supports the idea that mutations in *ahrC* and parallel pathways (such as chromosomal and ACME arginine-deiminase systems, *spoVG*) are selected in *S. aureus*, potentially widely in clinical settings. Moreover, our finding that mutations in *ahrC*/ACME do not adversely affect virulence during acute abscess infection in mice (Fig. 7H) suggests that mutation of arginine metabolic genes could be an evolutionary pathway to a colonization fitness produced without associated costs in virulence, possibly even with a gain.

Mutations in *ahrC* likely promote catabolism of arginine by *arcC* and *nos* pathways, improving, respectively, redox balance and energy production in the oxygen-limited gut environment. Typically, mutations in *ahrC*/ACME and *walKR* follow mutations in promoter/regulator regions in the glycolytic pathway that increase glycolytic flux (Fig. 2C). Thus, evolved mutants transport and metabolize the most desired nutrient (glucose) first. However, despite high glycolytic demand during infection (26), evolved *glcB sdaA* mutations that support fitness in the GI tract attenuated virulence during infection (Fig. 7). Infection likely involves fluctuating nutrient availabilities that require the metabolic versatility afforded by gene regulation. Increased metabolism of glucose due to the mutations might therefore represent a slash-and-burn strategy of competition for resources in which maximum growth potential is achieved through inactivation of important regulatory functions that are necessary for MRSA to occupy variety of niches.

Prior studies indicate a complex and largely inverse relationship between fitness for antimicrobial resistance and virulence (56, 57). Our studies reveal mechanisms and principles driving reciprocal interactions between these key traits, namely a previously unappreciated link with colonization and commensal microbiota (Fig. S6B). Broadly speaking, these data lead to a two-part framework that can help explain why phenotypic diversity is enriched in isolates from infecting but not from colonizing sites within natural populations of *S. aureus* (58). First, negative correlation among the metabolic requirements for colonization, virulence, and resistance underscores the wide range of phenotypes manifest during within-host adaptation to distinct disease states—a phenomenon known as pathoadaptation. The variability in how mutations impact these phenotypes as environmental conditions and microbiota change renders unpredictable the short-term advantages of pathoadaptive mutations and their long-term fitness, thereby complicating efforts to personalize treatment to each patient’s infection-adapted strain. For example, pathoadaptive mutations in the global regulator *agr* can both decrease virulence and colonization (16) while increasing antimicrobial tolerance (59). Consequently, the use of *agr* inhibitors, which are under development for treatment of MRSA infection, may be ill advised in certain patients (60). We expect that a better understanding of the tradeoffs involved in colonization, virulence, and resistance will enable more effective and syndrome-specific targeting of intervention strategies.

Second, commensal microbiota constrains adaptation of MRSA, suggesting an unappreciated role in maintaining wild-type metabolic and pathogenic flexibility necessary for long-term circulation of *S. aureus* through various niches in different hosts. This finding can help explain the ecology of new pathoadaptive variants. For example, VISA, and more generally antimicrobial-resistant hospital-associated MRSA clades, are only rarely encountered beyond vulnerable individuals within hospitals (61). In contrast, in healthy individuals outside hospitals, the barrier to colonization is likely higher than in hospitalized patients, where antimicrobial disruption of microbiota facilitates the spread of resistance-adapted MRSA strains with compromised fitness. This scenario highlights the potential drawbacks of prolonged, broad-spectrum empirical antimicrobial therapy and supports calls for microbiota transplantation as a strategy to decrease colonization by antimicrobial-resistant pathogens (62).

## METHODS

### Method Details

The Supplementary Materials provide detailed methods for constructing *S. aureus* strains and their growth conditions. Included are comprehensive descriptions of assays for intestinal colonization, skin infection, cytotoxicity, autolysis, and biofilm formation. Also described are methods for measuring antimicrobial susceptibility, mutation frequency, and performing genome sequencing, metagenomic, metabolomic, and statistical analyses. The strains and primers used in this study are listed in Tables S2 and S3, respectively.

## ACKNOWLEDGEMENTS

We thank members of the Shopsin, Torres, and Cadwell laboratories and Drs. Joel G. Belasco and Andrew Darwin for insightful discussions and critical comments on the manuscript, respectively. We thank Margie Alva, Juan Carrasquillo, and David Basnight for their help in the NYU Gnotobiotic Facility. We thank the NYU Langone Genome Technology Center (GTC))(RRID: SCR_017929) for expert support and NYU Langone Health Metabolomics Laboratory (RRID: SCR_017935) for its help in acquiring and analyzing the data presented. pIMAY∗ was a gift from Dr. Angelika Grundling (Addgene plasmid # 121441; http://n2t.net/addgene:121441; RRID: Addgene_121441). Graphical abstract, Fig. S2A, and Fig. S4 were created with BioRender.com.

This work was supported in part by the National Institute of Allergy and Infectious Diseases award numbers R01s AI140754 to B.S., V.J.T., and K.C.; AI137336 to B.S. and V.J.T.; K08 AI163457 to R.J.U., and funds from the NYU Langone Health Antimicrobial-Resistant Pathogens Program to B.S., A.P., and V.J.T. The GTC is partially supported by the Isaac Perlmutter Cancer Center Support grant P30CA016087 from the National Cancer Institute. The work reported in this paper was also supported by the Office of Research Infrastructure of the National Institutes of Health (NIH) under award numbers S10OD018522 and S10OD026880. J.S. reports relevant funding from the National Institutes of Health (NIH) grants DP2 AI164318-01.

## CONFLICTS OF INTERESTS

B.S. has consulted for Basilea Pharmaceutica. V.J.T. has received honoraria from Pfizer and MedImmune and is an inventor on patents and patent applications filed by New York University, which are currently under commercial license to Janssen Biotech Inc. Janssen Biotech Inc. provides research funding and other payments associated with a licensing agreement. K.C. has received research support from Pfizer, Takeda, Pacific Biosciences, Genentech, and AbbVie, consulted for or received honoraria from Vedanta, Genentech, and AbbVie, and is an inventor on US patent 10,722,600 and provisional patents 62/935,035 and 63/157,225. J.S. holds equity in Postbiotics Plus Research, has filed intellectual property applications related to the microbiome (reference numbers #63/299,607), and is on an advisory board and holds equity of Jona Health.

## REFERENCES

1. Antimicrobial Resistance C. 2022. Global burden of bacterial antimicrobial resistance in 2019: a systematic analysis. Lancet 399:629–655.

2. von Eiff C, Becker K, Machka K, Stammer H, Peters G. 2001. Nasal carriage as a source of *Staphylococcus aureus* bacteremia. Study Group. N Engl J Med 344:11–6.

3. Wertheim HF, Melles DC, Vos MC, van Leeuwen W, van Belkum A, Verbrugh HA, Nouwen JL. 2005. The role of nasal carriage in *Staphylococcus aureus* infections. Lancet Infect Dis 5:751–62.

4. Faden H, Lesse AJ, Trask J, Hill JA, Hess DJ, Dryja D, Lee YH. 2010. Importance of colonization site in the current epidemic of staphylococcal skin abscesses. Pediatrics 125:e618–24.

5. Kumar N, David MZ, Boyle-Vavra S, Sieth J, Daum RS. 2015. High *Staphylococcus aureus* colonization prevalence among patients with skin and soft tissue infections and controls in an urban emergency department. J Clin Microbiol 53:810–5.

6. Copin R, Sause WE, Fulmer Y, Balasubramanian D, Dyzenhaus S, Ahmed JM, Kumar K, Lees J, Stachel A, Fisher JC, Drlica K, Phillips M, Weiser JN, Planet PJ, Uhlemann AC, Altman DR, Sebra R, van Bakel H, Lighter J, Torres VJ, Shopsin B. 2019. Sequential evolution of virulence and resistance during clonal spread of community-acquired methicillin-resistant *Staphylococcus aureus*. Proc Natl Acad Sci U S A doi:10.1073/pnas.1814265116.

7. Buffie CG, Pamer EG. 2013. Microbiota-mediated colonization resistance against intestinal pathogens. Nat Rev Immunol 13:790–801.

8. Boyce JM, Havill NL. 2005. Nosocomial antibiotic-associated diarrhea associated with enterotoxin-producing strains of methicillin-resistant *Staphylococcus aureus*. Am J Gastroenterol 100:1828–34.

9. Boyce JM, Havill NL, Maria B. 2005. Frequency and possible infection control implications of gastrointestinal colonization with methicillin-resistant *Staphylococcus aureus*. J Clin Microbiol 43:5992–5.

10. Boyce JM, Havill NL, Otter JA, Adams NM. 2007. Widespread environmental contamination associated with patients with diarrhea and methicillin-resistant *Staphylococcus aureus* colonization of the gastrointestinal tract. Infect Control Hosp Epidemiol 28:1142–7.

11. Senn L, Clerc O, Zanetti G, Basset P, Prod’hom G, Gordon NC, Sheppard AE, Crook DW, James R, Thorpe HA, Feil EJ, Blanc DS. 2016. The stealthy superbug: the role of asymptomatic enteric carriage in maintaining a long-term hospital outbreak of ST228 methicillin-resistant *Staphylococcus aureus*. MBio 7:e02039–15.

12. Shopsin B, Kaveri SV, Bayry J. 2016. Tackling difficult *Staphylococcus aureus* infections: antibodies show the way. Cell Host Microbe 20:555–557.

13. Squier C, Rihs JD, Risa KJ, Sagnimeni A, Wagener MM, Stout J, Muder RR, Singh N. 2002. *Staphylococcus aureus* rectal carriage and its association with infections in patients in a surgical intensive care unit and a liver transplant unit. Infect Control Hosp Epidemiol 23:495–501.

14. Srinivasan A, Seifried SE, Zhu L, Srivastava DK, Perkins R, Shenep JL, Bankowski MJ, Hayden RT. 2010. Increasing prevalence of nasal and rectal colonization with methicillin-resistant *Staphylococcus aureus* in children with cancer. Pediatr Blood Cancer 55:1317–22.

15. Piewngam P, Khongthong S, Roekngam N, Theapparat Y, Sunpaweravong S, Faroongsarng D, Otto M. 2023. Probiotic for pathogen-specific *Staphylococcus aureus* decolonisation in Thailand: a phase 2, double-blind, randomised, placebo-controlled trial. Lancet Microbe 4:e75–e83.

16. Piewngam P, Zheng Y, Nguyen TH, Dickey SW, Joo HS, Villaruz AE, Glose KA, Fisher EL, Hunt RL, Li B, Chiou J, Pharkjaksu S, Khongthong S, Cheung GYC, Kiratisin P, Otto M. 2018. Pathogen elimination by probiotic *Bacillus* via signalling interference. Nature 562:532–537.

17. Leonidas Cardoso L, Durao P, Amicone M, Gordo I. 2020. Dysbiosis individualizes the fitness effect of antibiotic resistance in the mammalian gut. Nat Ecol Evol 4:1268–1278.

18. Boles BR, Thoendel M, Roth AJ, Horswill AR. 2010. Identification of genes involved in polysaccharide-independent *Staphylococcus aureus* biofilm formation. PLoS One 5:e10146.

19. Misawa Y, Kelley KA, Wang X, Wang L, Park WB, Birtel J, Saslowsky D, Lee JC. 2015. *Staphylococcus aureus* colonization of the mouse gastrointestinal tract Is modulated by wall teichoic acid, capsule, and surface proteins. PLoS Pathog 11:e1005061.

20. Elena SF, Lenski RE. 2003. Evolution experiments with microorganisms: the dynamics and genetic bases of adaptation. Nat Rev Genet 4:457–69.

21. Canfield GS, Schwingel JM, Foley MH, Vore KL, Boonanantanasarn K, Gill AL, Sutton MD, Gill SR. 2013. Evolution in fast forward: a potential role for mutators in accelerating *Staphylococcus aureus* pathoadaptation. J Bacteriol 195:615–28.

22. Diep BA, Gill SR, Chang RF, Phan TH, Chen JH, Davidson MG, Lin F, Lin J, Carleton HA, Mongodin EF, Sensabaugh GF, Perdreau-Remington F. 2006. Complete genome sequence of USA300, an epidemic clone of community-acquired meticillin-resistant *Staphylococcus aureus*. Lancet 367:731–9.

23. Mwangi MM, Wu SW, Zhou Y, Sieradzki K, de Lencastre H, Richardson P, Bruce D, Rubin E, Myers E, Siggia ED, Tomasz A. 2007. Tracking the *in vivo* evolution of multidrug resistance in *Staphylococcus aureus* by whole-genome sequencing. Proceedings of the National Academy of Sciences 104:9451–9456.

24. Rochat T, Bohn C, Morvan C, Le Lam TN, Razvi F, Pain A, Toffano-Nioche C, Ponien P, Jacq A, Jacquet E, Fey PD, Gautheret D, Bouloc P. 2018. The conserved regulatory RNA RsaE down-regulates the arginine degradation pathway in *Staphylococcus aureus*. Nucleic Acids Res 46:8803–8816.

25. Brugiroux S, Beutler M, Pfann C, Garzetti D, Ruscheweyh HJ, Ring D, Diehl M, Herp S, Lotscher Y, Hussain S, Bunk B, Pukall R, Huson DH, Munch PC, McHardy AC, McCoy KD, Macpherson AJ, Loy A, Clavel T, Berry D, Stecher B. 2016. Genome-guided design of a defined mouse microbiota that confers colonization resistance against *Salmonella enterica* serovar Typhimurium. Nat Microbiol 2:16215.

26. Vitko NP, Grosser MR, Khatri D, Lance TR, Richardson AR. 2016. Expanded glucose import capability affords *Staphylococcus aureus* optimized glycolytic flux during Infection. mBio 7.

27. Knezevic I, Bachem S, Sickmann A, Meyer HE, Stulke J, Hengstenberg W. 2000. Regulation of the glucose-specific phosphotransferase system (PTS) of *Staphylococcus carnosus* by the antiterminator protein GlcT. Microbiology (Reading) 146 (Pt 9):2333–2342.

28. Langbein I, Bachem S, Stulke J. 1999. Specific interaction of the RNA-binding domain of the *Bacillus subtilis* transcriptional antiterminator GlcT with its RNA target, RAT. J Mol Biol 293:795–805.

29. Greenwich J, Reverdy A, Gozzi K, Di Cecco G, Tashjian T, Godoy-Carter V, Chai Y. 2019. A Decrease in serine levels during growth transition triggers biofilm formation in *Bacillus subtilis*. J Bacteriol 201.

30. Pizer LI, Potochny ML. 1964. Nutritional and regulatory aspects of serine metabolism in *Escherichia Coli*. J Bacteriol 88:611–9.

31. Kitamoto S, Alteri CJ, Rodrigues M, Nagao-Kitamoto H, Sugihara K, Himpsl SD, Bazzi M, Miyoshi M, Nishioka T, Hayashi A, Morhardt TL, Kuffa P, Grasberger H, El-Zaatari M, Bishu S, Ishii C, Hirayama A, Eaton KA, Dogan B, Simpson KW, Inohara N, Mobley HLT, Kao JY, Fukuda S, Barnich N, Kamada N. 2020. Dietary L-serine confers a competitive fitness advantage to *Enterobacteriaceae* in the inflamed gut. Nat Microbiol 5:116–125.

32. Makhlin J, Kofman T, Borovok I, Kohler C, Engelmann S, Cohen G, Aharonowitz Y. 2007. *Staphylococcus aureus* ArcR controls expression of the arginine deiminase operon. J Bacteriol 189:5976–86.

33. Reslane I, Halsey CR, Stastny A, Cabrera BJ, Ahn J, Shinde D, Galac MR, Sladek MF, Razvi F, Lehman MK, Bayles KW, Thomas VC, Handke LD, Fey PD. 2022. Catabolic Ornithine carbamoyltransferase activity facilitates growth of *Staphylococcus aureus* in defined medium lacking glucose and arginine. mBio 13:e0039522.

34. Lindgren JK, Thomas VC, Olson ME, Chaudhari SS, Nuxoll AS, Schaeffer CR, Lindgren KE, Jones J, Zimmerman MC, Dunman PM, Bayles KW, Fey PD. 2014. Arginine deiminase in *Staphylococcus epidermidis* functions to augment biofilm maturation through pH homeostasis. J Bacteriol 196:2277–89.

35. Kinkel TL, Ramos-Montanez S, Pando JM, Tadeo DV, Strom EN, Libby SJ, Fang FC. 2016. An essential role for bacterial nitric oxide synthase in *Staphylococcus aureus* electron transfer and colonization. Nat Microbiol 2:16224.

36. Fey PD, Endres JL, Yajjala VK, Widhelm TJ, Boissy RJ, Bose JL, Bayles KW. 2013. A genetic resource for rapid and comprehensive phenotype screening of nonessential *Staphylococcus aureus* genes. MBio 4:e00537–12.

37. Caballero-Flores G, Pickard JM, Fukuda S, Inohara N, Nunez G. 2020. An enteric pathogen subverts colonization resistance by evading competition for amino acids in the gut. Cell Host Microbe 28:526–533 e5.

38. Sasabe J, Miyoshi Y, Rakoff-Nahoum S, Zhang T, Mita M, Davis BM, Hamase K, Waldor MK. 2016. Interplay between microbial d-amino acids and host d-amino acid oxidase modifies murine mucosal defence and gut microbiota. Nat Microbiol 1:16125.

39. Howden BP, McEvoy CR, Allen DL, Chua K, Gao W, Harrison PF, Bell J, Coombs G, Bennett-Wood V, Porter JL, Robins-Browne R, Davies JK, Seemann T, Stinear TP. 2011. Evolution of multidrug resistance during *Staphylococcus aureus* infection involves mutation of the essential two component regulator WalKR. PLoS Pathog 7:e1002359.

40. Hafer C, Lin Y, Kornblum J, Lowy FD, Uhlemann AC. 2012. Contribution of selected gene mutations to resistance in clinical isolates of vancomycin-intermediate *Staphylococcus aureus*. Antimicrob Agents Chemother 56:5845–51.

41. Monk IR, Shaikh N, Begg SL, Gajdiss M, Sharkey LKR, Lee JYH, Pidot SJ, Seemann T, Kuiper M, Winnen B, Hvorup R, Collins BM, Bierbaum G, Udagedara SR, Morey JR, Pulyani N, Howden BP, Maher MJ, McDevitt CA, King GF, Stinear TP. 2019. Zinc-binding to the cytoplasmic PAS domain regulates the essential WalK histidine kinase of *Staphylococcus aureus*. Nat Commun 10:3067.

42. Mwangi MM, Wu SW, Zhou Y, Sieradzki K, de Lencastre H, Richardson P, Bruce D, Rubin E, Myers E, Siggia ED, Tomasz A. 2007. Tracking the *in vivo* evolution of multidrug resistance in *Staphylococcus aureus* by whole-genome sequencing. Proc Natl Acad Sci U S A 104:9451–6.

43. Shoji M, Cui L, Iizuka R, Komoto A, Neoh HM, Watanabe Y, Hishinuma T, Hiramatsu K. 2011. *walK* and *clpP* mutations confer reduced vancomycin susceptibility in *Staphylococcus aureus*. Antimicrob Agents Chemother 55:3870–81.

44. Laabei M, Uhlemann AC, Lowy FD, Austin ED, Yokoyama M, Ouadi K, Feil E, Thorpe HA, Williams B, Perkins M, Peacock SJ, Clarke SR, Dordel J, Holden M, Votintseva AA, Bowden R, Crook DW, Young BC, Wilson DJ, Recker M, Massey RC. 2015. Evolutionary trade-offs underlie the multi-faceted virulence of *Staphylococcus aureus*. PLoS Biol 13:e1002229.

45. Rose HR, Holzman RS, Altman DR, Smyth DS, Wasserman GA, Kafer JM, Wible M, Mendes RE, Torres VJ, Shopsin B. 2015. Cytotoxic virulence predicts mortality in nosocomial pneumonia due to methicillin-resistant *Staphylococcus aureus*. J Infect Dis 211:1862–74.

46. Voyich JM, Braughton KR, Sturdevant DE, Whitney AR, Said-Salim B, Porcella SF, Long RD, Dorward DW, Gardner DJ, Kreiswirth BN, Musser JM, DeLeo FR. 2005. Insights into mechanisms used by *Staphylococcus aureus* to avoid destruction by human neutrophils. J Immunol 175:3907–19.

47. Liu X, Zhang S, Sun B. 2016. SpoVG regulates cell wall metabolism and oxacillin resistance in methicillin-resistant *Staphylococcus aureus* strain N315. Antimicrob Agents Chemother 60:3455–61.

48. Miller WR, Bayer AS, Arias CA. 2016. Mechanism of action and resistance to daptomycin in *Staphylococcus aureus* and enterococci. Cold Spring Harb Perspect Med 6.

49. Cui L, Isii T, Fukuda M, Ochiai T, Neoh HM, Camargo IL, Watanabe Y, Shoji M, Hishinuma T, Hiramatsu K. 2010. An RpoB mutation confers dual heteroresistance to daptomycin and vancomycin in *Staphylococcus aureus*. Antimicrob Agents Chemother 54:5222–33.

50. Wang R, Braughton KR, Kretschmer D, Bach T-HL, Queck SY, Li M, Kennedy AD, Dorward DW, Klebanoff SJ, Peschel A, DeLeo FR, Otto M. 2007. Identification of novel cytolytic peptides as key virulence determinants for community-associated MRSA. Nat Med 13:1510–1514.

51. Li M, Cheung GY, Hu J, Wang D, Joo HS, Deleo FR, Otto M. 2010. Comparative analysis of virulence and toxin expression of global community-associated methicillin-resistant *Staphylococcus aureus* strains. J Infect Dis 202:1866–76.

52. Yan J, Liao C, Taylor BP, Fontana E, Amoretti LA, Wright RJ, Littmann ER, Dai A, Waters N, Peled JU, Taur Y, Perales M-A, Siranosian BA, Bhatt AS, van den Brink MRM, Pamer EG, Schluter J, Xavier JB. 2022. A compilation of fecal microbiome shotgun metagenomics from hematopoietic cell transplantation patients. Scientific Data 9:219.

53. Schluter J, Djukovic A, Taylor BP, Yan J, Duan C, Hussey GA, Liao C, Sharma S, Fontana E, Amoretti LA, Wright RJ, Dai A, Peled JU, Taur Y, Perales M-A, Siranosian BA, Bhatt AS, van den Brink MRM, Pamer EG, Xavier JB. 2023. The TaxUMAP atlas: Efficient display of large clinical microbiome data reveals ecological competition in protection against bacteremia. Cell Host & Microbe 31:1126–1139.e6.

54. Reslane I, Handke LD, Watson GF, Shinde D, Ahn JS, Endres JL, Razvi F, Gilbert EA, Bayles KW, Thomas VC, Lehman MK, Fey PD. 2024. Glutamate-dependent arginine biosynthesis requires the inactivation of *spoVG*, *sarA*, and *ahrC* in *Staphylococcus aureus*. J Bacteriol 206:e0033723.

55. Ng PC, Henikoff S. 2001. Predicting deleterious amino acid substitutions. Genome Res 11:863–74.

56. Andersson DI. 2006. The biological cost of mutational antibiotic resistance: any practical conclusions? Curr Opin Microbiol 9:461–5.

57. Andersson DI, Hughes D. 2010. Antibiotic resistance and its cost: is it possible to reverse resistance? Nat Rev Microbiol 8:260–71.

58. Young BC, Wu CH, Gordon NC, Cole K, Price JR, Liu E, Sheppard AE, Perera S, Charlesworth J, Golubchik T, Iqbal Z, Bowden R, Massey RC, Paul J, Crook DW, Peto TE, Walker AS, Llewelyn MJ, Wyllie DH, Wilson DJ. 2017. Severe infections emerge from commensal bacteria by adaptive evolution. Elife 6:e30637.

59. Kumar K, Chen J, Drlica K, Shopsin B. 2017. Tuning of the lethal response to multiple stressors with a single-site mutation during clinical infection by *Staphylococcus aureus*. MBio 8:e01476–17.

60. Khan BA, Yeh AJ, Cheung GY, Otto M. 2015. Investigational therapies targeting quorum-sensing for the treatment of *Staphylococcus aureus* infections. Expert Opin Investig Drugs 24:689–704.

61. Howden BP, Davies JK, Johnson PD, Stinear TP, Grayson ML. 2010. Reduced vancomycin susceptibility in *Staphylococcus aureus*, including vancomycin-intermediate and heterogeneous vancomycin-intermediate strains: resistance mechanisms, laboratory detection, and clinical implications. Clin Microbiol Rev 23:99–139.

62. Woodworth MH, Conrad RE, Haldopoulos M, Pouch SM, Babiker A, Mehta AK, Sitchenko KL, Wang CH, Strudwick A, Ingersoll JM, Philippe C, Lohsen S, Kocaman K, Lindner BG, Hatt JK, Jones RM, Miller C, Neish AS, Friedman-Moraco R, Karadkhele G, Liu KH, Jones DP, Mehta CC, Ziegler TR, Weiss DS, Larsen CP, Konstantinidis KT, Kraft CS. 2023. Fecal microbiota transplantation promotes reduction of antimicrobial resistance by strain replacement. Sci Transl Med 15:eabo2750.

