## Supplemental Figure S1-S6, Supplemental Tables S1-S3 for "Microbiota and metabolic adaptation shape *Staphylococcus aureus* virulence and antimicrobial resistance during intestinal colonization"

**CONTENTS**

**Supplementary Text (Methods)**

**Supplementary Figures (S1-S6)**

Fig. S1. Evolved mutants are not hypermutable

Fig. S2. Mutations in selected genes identified during intestinal adaptation of *S. aureus* in germ-free mice

Fig. S3. Bacterial burden and cadmium resistance marker for competition assays of *S. aureus* colonization fitness

Fig. S4. Evolved mutations in *glcB* and *glcT* are predicted to increase glucose import

Fig. S5. *glcB* *walK* mutant demonstrates increased fitness for colonization throughout the gut, and evolved changes have limited impact on cytotoxicity compared to parental strain LAC

Fig. S6. Models of adaptation during intestinal colonization

**Supplementary Tables**

Table S1. List of LAC evolved strains used in cefoxitin and daptomycin Etests. Related to Fig. 7C-7E.

Table S2. List of bacterial strains used in this study.

Table S3. List of oligonucleotides used in this study.

**Supplementary Datasets (S1-S5)**

Dataset S1. Matrix of single nucleotide variants (SNVs) in 201 colonies of strain LAC isolated from stool following evolution in germ-free mice housed in 4 cages for 5 weeks. SNP were not identified in 1 isolate, which is not included in the table.

Dataset S2. Matrix of single nucleotide variants (SNVs) in 92 colonies of strain JH1 isolated from stool following evolution in germ-free mice for 5 weeks. SNP were not identified in 26 isolates, which are not included in the table.

Dataset S3. SNVs in 20 colonies of strain LAC isolated from stool following evolution in germ-free mice housed 4 cages (5 colonies per cage) for 1 week.

Dataset S4. SNVs in clonal MRSA isolates (BS1208 and BS1210) recovered sequentially from the bloodstream of a patient treated with vancomycin. After extensive therapy, the bacterium developed vancomycin heteroresistance.

Dataset S5. Evolved mutations in *S. aureus* in human gut. SNV analysis was performed in coding sequences and promoter regions from human gut metagenomic samples with focus on pathways that were involved in murine gut adaptation.

**Supplementary References**

**Supplementary Text (Materials and methods)**

***Bacterial Strains and Growth Conditions***

Bacterial cells were grown in tryptic soy broth (TSB) at 37ºC with rotary shaking at 180 rpm. Colony formation was on tryptic soy agar (TSA) or TSA with 5% sheep blood (Becton Dickenson, Franklin Lakes, NJ), incubated at 37ºC. *E. coli* IM08B (1), which was used for cloning, was grown in Luria-Bertani broth medium. Appropriate antibiotics (chloramphenicol to 10 μg/mL or 20 μg/mL, ampicillin 100 μg/mL, erythromycin to 5 μg/mL or 10 μg/mL, and tetracycline to 4 μg/mL) were used as needed.

For analysis of *in vitro* growth curves, overnight cultures grown in TSB were washed in PBS, cell density was normalized by OD_600_, and cultures were diluted to the same starting OD_600_ in 150 μL culture media in chemically defined medium (CDM) (2). Growth was monitored at 37°C using Honeycomb Microplates (Oy Growth Curves Ab Ltd, Finland) (150 µL/well) and a Bioscreen C (Oy Growth Curves Ab Ltd, Finland) to measure OD_600_ at 15 or 30-min intervals. CDM, prepared as previously described (3), lacked glucose unless otherwise stated. The curves represent averaged values from three biological replicates.

***Intestinal colonization of mice***

For LAC evolution experiments, germ-free C57BL/6J mice were bred on site to generate animals for experimentation in the New York University (NYU) School of Medicine Gnotobiotics Animal Facility (4). Sixteen age and gender-matched, 8–10-week-old mice were used. Absence of fecal bacteria and fungi was confirmed by aerobic culture in brain heart infusion, Sabouraud, and nutrient broth (Sigma-Aldrich, St. Louis, MO) as well as qPCR for bacterial 16S and eukaryotic 18S rRNA genes through sampling of stool from individual cages in each isolator on a monthly basis. Mice were transferred into individually ventilated TISOcages (Tecniplast, West Chester, PA) for colonization studies to maintain sterility under positive air pressure. Mice were orally gavaged with ~5 × 10^8^ cfu of strain LAC. Stool samples were collected from each mouse weekly for 10 weeks after inoculation with LAC. Screw-cap tubes (2 mL) filled with 1.0 mm beads were weighed before and after the addition of stool to determine stool weight. Sterile phosphate-buffered saline (PBS) (1 mL) was added to each tube, which were vigorously shaken in a bead-beater (MP Biomedicals, Santa Ana, CA) for 60 s. Colony formation was on TSA with 5% sheep blood (Becton Dickenson, Franklin Lakes, NJ), incubated at 37ºC, and morphologically distinct colonies were subcultured and stocked at -80°C. JH1 evolution experiments in germ-free mice, performed as described for LAC, involved 14 germ-free C57BL/6 mice, ~3 × 10^8^ cfu of JH1 inoculum, and three ISOcages.

Conventional mice were screened for *S. aureus* colonization prior to inoculation with LAC by plating stool on CHROMagar Staph aureus (CHROMagar, Paris, France). For evolution experiments, seven C57BL/6J mice (eight weeks old, males and females) were orally gavaged with approximately 1.5 × 10^8^ cfu of LAC. Stool samples were collected weekly for five weeks after inoculation with LAC, and LAC burden was determined by plating stool samples on BBL CHROMagar MRSA II (Becton Dickenson, Franklin Lakes, NJ).

Minimal defined flora mice were colonized by a consortium of 15 bacterial strains representing the murine gut microbiota (Oligo-MM^12^+FA^3^) (5), as described (6). Evolution experiments were performed as described for germ-free mice. Stool samples were collected weekly for five weeks after inoculation with LAC, and LAC burden was determined by plating stool samples on BBL CHROMagar MRSA II (Becton Dickenson, Franklin Lakes, NJ). Mice containing the consortium were kept on a standard, germ-free (autoclaved) diet; thus, the absence of adaptive mutations in the presence of microbiota does not require changes in diet.

*Competition assays.* For in *vivo* competition experiments in germ-free or conventional mice wild-type reference strain LAC was grown separately from the evolved competitor strain overnight at 37 °C, diluted 100-fold, subcultured for 4 h, and mixed at a 1:1 ratio of ~1 × 10^8^ cfu prior to inoculation. One of the two competitors contained a chromosomally integrated cadmium resistance marker (SaPI1 *att*C::*cadCA*) to distinguish the strains following plating of serial dilutions for bacterial burden (cfu); relative numbers of cfu in feces were determined by plating on nonselective and selective plates to select resistant strains. For germ-free mice, homogenized stool samples were plated on TSA with and without 0.3 mM cadmium chloride (Hampton Research, Aliso Viejo, CA). For conventional mice, homogenized stool samples were plated on CHROMagar Staph aureus (CHROMagar, Paris, France) with and without 0.3 mM cadmium chloride (Hampton Research). Output of total and cadmium-resistant cfu from stool were determined, and the competitive index was calculated as the ratio of the two strains in competition.

*Intestinal localization*. Intestinal tissue mucus, cecal content, and stool were collected from each mouse on day 6 post-inoculation of germ-free mice. Small intestine (5 cm proximal to the cecum), cecum, and colon (5 cm distal to the cecum) luminal contents were removed, and tissues were rinsed thoroughly three times with sterile PBS using 5-mL syringes (BD) and 20-G animal feeding needles (Pet Surgical, Phoenix, AZ). Tissue mucus was harvested by scraping and suspended in 1 mL sterile PBS to be used for serial dilution and bacterial burden enumeration, as described (7). Approximately 10 μL of cecal content was added to 1 mL sterile PBS for bacterial burden enumeration.

***Skin infection of mice***

For the skin infections, overnight cultures of the indicated strain were diluted 1:100 into fresh TSB medium, grown to exponential phase (3.5 h at 37°C shaking at 180 rpm), washed in PBS and normalized to ~1 × 10^8^ cfu. Five-week-old female ND4 Swiss Webster mice (Envigo, Inc., Indianapolis, IN) were anesthetized intraperitoneally with 300 μL of Avertin (2,2,2-tribromoethanol dissolved in tert-Amyl alcohol and diluted to a final concentration of 2.5% [vol/vol] in sterile PBS) and the back/flank region was shaved with mechanical clippers. Appoximately 100 μL of bacteria, resulting in a ~1 x 10^7^ cfu inoculum was injected subcutaneously into both hind flanks of shaved mice (8). Every 24 hours, mice were briefly anesthetized with inhaled isoflurane, and lesions were digitally imaged (Apple Inc., Cupertino, CA). Abscess area was measured using ImageJ software (9). To assess bacterial burden, 8-mm punch biopsy (Integra Life Sciences, Princeton, NJ) samples were obtained from mouse lesions at 72 h postinfection, and the tissues were homogenized in 2 mL conical screw cap tubes (Fisher Scientific) with 1 mL sterile PBS and a single 0.25” ceramic sphere (MP Biomedicals, Santa Ana, CA) for three cycles in a bead beater (FastPrep-24, MP Biomedicals, Santa Ana, CA) for 60 seconds at 4 m/s. Homogenized abscess tissues were serially diluted and enumerated on TSA (10).

***Construction of mutants***

Transposon mutants were generated by transducing *Bursa aurealis* insertions, obtained from the University of Nebraska transposon mutant (ΦNE) library (11), into LAC or evolved mutant strains using phages 80α and Φ11. Cadmium-resistance marked stains were generated by transduction from strain VJT32.58 (LAC CdR) (12) using phage 80α or Φ11.

Strain LAC WalK^H271Y^ was generated via allelic exchange. The DNA fragment containing WalK^H271Y^ mutation, a silent PstI site (*walK* nucleotide 1302 A to G) (13), and flanking regions of vector plasmid pIMAY* (14) containing the multiple cloning site was synthesized as a gBlock (IDT, Coralville, IA). Gibson assembly of the WalK^H271Y^ fragment and pIMAY* vector was performed using NEBuilder® HiFi DNA Assembly Master Mix (New England Biolabs, Ipswich, MA). The resulting plasmid pCZ1 was transformed into *Escherichia coli* strain IM08B (1), and primer E239K_scr_F was used to confirm the desired mutation in pCZ1 before electroporation into LAC. Allelic exchange was performed as previously described (15). *walK* PCR products, amplified with primers walRK_F2 and walRK_R1, were digested with PstI-HF (New England Biolabs, Ipswich, MA) to identify double recombinant colonies that contain the desired construct. Primer E239K_scr_F and Sanger sequencing was used to confirm the presence of the WalK^H271Y^ mutation.

***Real-time qRT-PCR assays***

Overnight cultures grown in TSB were washed with PBS, diluted (OD_600_ of 0.05), and inoculated in 250 mL flasks (PYREX) with 25 mL CDM 1% casamino acids without glucose or 20 mL CDM 1% casamino acids with 14 mM glucose. Cells were grown to late-exponential phase (~4 h) and concentrated by centrifugation for 10 min at 12,000 g. Cells were resuspended with 1 mL TRIzol (Invitrogen, Waltham, MA) and added to Lysing Matrix B (MP Biomedicals, Santa Ana, CA) for bead-beating 3 times at 6 m/s for 30 seconds (FastPrep-24, MP Biomedicals, Santa Ana, CA). Cell lysates were centrifuged for 10 min at 12,000 g. The upper phase was transferred into 500 μL TRIzol and incubated at room temperature for 5 min. Cell lystates were extracted with chloroform (200μL), and samples of the aqueous phase were transferred to fresh tubes containing 600 μL isopropanol to precipitate RNA. An RNAeasy Mini Kit (Qiagen, Hilden, Germany) was used to further purify RNA. RNA (100-500 ng per sample depending on the abundance of the target gene), treated with TURBO DNA-free kit (Invitrogen), was used to synthesize cDNAs using a SuperScript™ III First-Strand Synthesis System (Invitrogen). Real-time reverse transcription quantitative PCR (qRT-PCR) was performed using QuantiNova™ SYBR Green PCR Kit (Qiagen, Hilden, Germany) in a CFX96 Real-Time System (Bio-Rad, Hercules, CA). Primers (*glcB*, ptsG_F3 and ptsG_R3; 16S rRNA, 16S_rRNA_F and 16S_rRNA_R) (16) were synthesized by IDT Inc. (Coralville, IA). The 2^–∆∆Ct^ method was used to calculate the relative fold gene expression (17), and mRNA levels of the target gene in each strain were normalized to the corresponding gene in the LAC strain.

***Autolysis assays***

Overnight cultures were washed with PBS, normalized to OD_600_ of 2 in two tubes each with 1 mL PBS, pelleted, and resuspended in 1 mL PBS or 1 mL PBS+0.1% Triton X-100 (Sigma-Aldrich, St. Louis, MO). Aliquots (100 μL) were added to Honeycomb Microplates (Oy Growth Curves Ab Ltd, Finland), and OD_600_ was measured every 30 min using a Bioscreen C microplate reader (Oy Growth Curves Ab Ltd, Finland). Percent absorbance was calculated relative to initial OD_600_.

***Biofilm formation assays***

Biofilm was assayed using a protocol adapted from a previous example (18). Briefly, overnight cultures were diluted 1:100 in TSB supplemented with 0.25% w/v D-(+)-glucose. Aliquots of 200 μL were transferred to a flat-bottom tissue culture-treated 96-well microtiter plate (Corning, Corning, NY). After static incubation at 37 °C for 24 h, the supernatant was removed and wells were washed twice with sterile PBS. To stain the biofilm, 200 μL of 0.1% (w/v) crystal violet was added to the wells and removed after 15 min of incubation. Excess crystal violet was washed with water. After adding 33% (v/v) acetic acid, biofilm was quantified by measuring absorbance at 595 nm with a BioTek Synergy Neo2 plate reader (Agilent, Santa Clara, CA).

***Antimicrobial Susceptibility Testing***

Minimum inhibition concentrations (MICs) for vancomycin, daptomycin, and cefoxitin were determined by Etest, according to manufacturer’s instructions (BioMérieux, Marcy-l'Étoile, France). Colonies, grown on TSA with 5% sheep blood plates (Becton Disckenson, Franklin Lakes, NJ), were suspended in 0.9% saline solution, adjusted for cell density using a 0.5 McFarland standard (~10^5^ cells), and spread evenly (200 μL) on Mueller-Hinton agar plates. Plates were incubated at 37°C for 24 h.

Heterogeneous resistance to glycopeptides was determined using vancomycin-teicoplanin double-sided gradient GRD Etest (BioMérieux, Marcy-l'Étoile, France) strips, as described (19). Cell suspensions, prepared and plated as for MIC determination, were plated and incubated at 37°C for 48 h. Results were interpreted by two independent observers.

Modified population analysis profiles (PAPs) were performed as described (20). Briefly, overnight cultures were serially diluted in phosphate-buffered saline (OD_600_ of 0.05) and plated (in 10 μL droplets) on a series of Brain Heart Infusion agar (BD) plates with the following concentrations of vancomycin (Sigma-Aldrich): 0, 0.5, 1, 1.5, 2, 3, 4, 6, and 8 μg/mL. Plates were incubated at 37°C for 48 h and bacterial cfu were plotted (log_10_cfu/mL) against vancomycin concentration. hVISA reference strain Mu3 (21) was used as a control.

***Spontaneous Rifampin-Resistant Mutant Recovery Assay***

Mutation frequency to rifampin was determined as previously described (22). Briefly, overnight cultures were grown for 16 h in TSB at 37°C with 180 rpm shaking. Samples were serially diluted and plated onto TSA plates with and without 40 ng/mL rifampin (Sigma-Aldrich, St. Louis, MO). After spot plating, plates were incubated at 37°C, and colonies were counted after 24 h.

***Cytotoxicity Assays***

Human polymorphonuclear neutrophils (hPMNs) were isolated as described previously (23) from Leukopaks obtained from deidentified donors from the New York Blood Center. *S. aureus* overnight cultures were subcultured in fresh TSB for 5 h at 37 °C. Culture supernatant obtained from early logarithmic-phase growth were used for differentiating cytotoxic activity. Briefly, overnight cultures were dilute 1:100 with fresh TSB, and the diluted culture was regrown for 6 h at 37 °C. Bacteria were pelleted, and filtered (0.2 μm) supernatants were diluted and added to 2 × 10^5^ hPMNs per well for a final volume of 100 μL per well in RPMI 1640 (Gibco, Billings, MT) supplemented with 10% FBS. hPMNs were intoxicated with the culture supernatant from the indicated strain for 2 h at 37 °C and 5% CO_2_. hPMN viability was measured using CellTiter 96 Aqueous One Solution (Promega, Madison, WI) Cell viability was measured by absorbance at 492 nm using an Envision 2103 Multilabel reader (PerkinElmer, Waltham, MA).

***Whole-genome sequencing and data analysis***

For LAC and LAC evolution experiments in germ-free mice, we genome sequenced the parental strain and 202 and 118 evolved strains, respectively, following isolation of single colonies from the stool of mice 5 weeks post-inoculation. DNA was extracted from each MRSA isolate as previously described (8), and sequence libraries were generated (Illumina, San Diego, CA) following the manufacturer’s recommendations. DNA was extracted using the KingFisher Flex Purification System (Thermo Fisher, Waltham, MA), and whole-genome sequencing was performed using Illumina NovaSeq 6000 (~450x coverage).

For sequencing pooled colony populations, stool from mice in all four cages was collected at weeks 1, 2, 3, 4, and 5 post-inoculation, pooled, and plated on TSA with 5% sheep blood (Becton Disckenson, Franklin Lakes, NJ). After growth overnight at 37°C, bacterial colonies were scraped from the agar plates, suspended in sterile PBS, and used for DNA extraction.

Paired-end Illumina reads were mapped using the RefSeq assembly GCF_015475575.1 (LAC; this reference assembly differed from our input strain at four positions that were accounted for) or GCF_000017125.1 (JH1; corrected with the sequencing reads from our JH1 input strain using pilon v. 1.24(24) and reannotated using PGAP v. 2021-11-29.build5742) (25). Sequencing reads were aligned to the respective reference assembly (above) with smalt v. 0.7.6 (26) using default parameters. Sorting and indexing of the alignment were done using samtools v. 1.9 (27). For purified colonies, single nucleotide variants (SNVs) were called using bcftools mpileup v. 1.9(27) and filtered according to established quality criteria (28); briefly, we required each SNV to have a quality score of at least 20 and to be covered at least 5x by at least one forward and one reverse read and alternate alleles to be supported by at least 75% of reads. For pooled populations, the population census of SNVs was determined by freebayes v. 1.3.5 (29) with the following flags: --legacy-gls --pooled-continuous -F 0.01 -C 1. Annotations were transferred from the respective reference assembly to the called SNVs using SnpEff v. 5.1 (30).

To obtain *S. aureus* FPR3757 (RefSeq GCF_000013465.1) locus tags for LAC and JH1, the predicted protein sequences for each pair of assemblies were matched using blastp (31) with an e-value threshold of 10^-9^. Only the top hit for each query was retained if the identity was greater than 95% and each of the two sequences in the alignment was covered by at least 80%. Locus tag equivalences for non-coding RNA sequences were established with blastn (31) in megablast mode using the same parameters as above.

For detecting the presence of ACME and *mecA* in our isolates, we identified the nucleotide sequences of *arcA* (SAUSA300_0065), *opp3A* (SAUSA300_0073), and *mecA* (SAUSA300_0032) from the *S. aureus* FPR3757 RefSeq assembly (GCF_000013465.1) and used them as a reference for ariba v. 2.14.6 (32). ACME was assumed to be present in an isolate if both *arcA* and *opp3A* were predicted to be present at more than 98% identity and more than 75% coverage; the same thresholds were used to determine *mecA* presence.

*Phylogenetic Analysis*. For maximum-likelihood phylogenies, a core genome alignment was compiled from passing SNV and invariant site calls (*Whole-Genome Sequencing and Comparative Genomics*). A phylogeny was obtained from the resulting alignment using RAxML v. 8.2.12 (33) with 1,000 rapid bootstrap replicates.

*Visualization****.*** Plots depicting the presence and absence of genetic elements were created using R v. 4.1.1 (34) and the ggplot and patchwork libraries. Phylogenies were plotted with iToL (35).

Whole genome sequencing data were deposited to NCBI BioProject PRJNA988476.

***RNA structure predictions***

The *S. aureus* putative *glcB* terminator and ribonucleic-antiterminator (RAT) sequences were predicted based on the *ptsG* terminator and RAT sequences of *Bacillus subtills.*(36) RNA structure predictions of the terminator loops were performed using the Mfold web server (37).

***Metabolomic analysis***

*Extraction of metabolites from mouse stool*. Fecal pellets homogenized in PBS (about 20 mg/mL) from three mice in cage 1 from pre- and one-week post-inoculation were used in the analysis. Prior to extraction, samples were moved from -80 °C storage to wet ice and thawed. Extraction buffer, consisting of 80% methanol (Fisher Scientific, City, State) and 500 nM metabolomics amino acid mix standard (Cambridge Isotope Laboratories, Inc., City, State), was prepared and placed on dry ice. For extraction, 50 µL of sample was mixed with 950 µL of extraction buffer in 2.0 mL screw cap vials containing ~100 µL of disruption beads (Research Products International, Mount Prospect, IL). Samples were lysed by repeated homogenization

(10 cycles of 30 sec homogenization time at 6 m/s followed by a 30-second pause) in a BeadBlaster (Benchmark Scientific, Edison, NJ). Cell debris was removed by centrifugation (21,000 g for 3 min at 4 °C) and the supernatant (450 µL) was transferred to a 1.5 mL tube, dried by speedvac (Thermo Fisher, Waltham, MA) and reconstituted in 50 µL of Optima LC/MS grade water (Fisher Scientific, Waltham, MA). Samples were sonicated for 2 mins, centrifuged at 21,000 g for 3 min at 4 °C. 20 µL, and transferred to liquid chromatography vials containing glass inserts for analysis. The remaining sample was placed in -80 °C for long-term storage.

*LC-MS/MS.* Samples were subjected to LC-MS analysis to detect and quantify known peaks. Metabolite extraction was carried out on each sample, as previously described (38). The LC column was a Millipore™ ZIC-pHILIC (2.1 x150 mm, 5 μm) coupled to a Dionex Ultimate 3000™ system (Thermo Scientific, Waltham, MA) and the column oven temperature was set to 25 °C for the gradient elution. A flow rate of 100 μL/min was used with the following buffers; A) 10 mM ammonium carbonate in water, pH 9.0, and B) neat acetonitrile. The gradient profile was as follows; 80-20% B (0-30 min), 20-80% B (30-31 min), 80-80% B (31-42 min). The injection volume was set to 2 μL for all analyses (42 min total run time per injection). MS analyses were carried out by coupling the LC system to a Thermo Q Exactive HF™ mass spectrometer operating in heated electrospray ionization mode (HESI). Method duration was 30 min with a polarity switching data-dependent Top 5 method for both positive and negative modes. The spray voltage for both positive and negative modes was 3.5kV and capillary temperature was set to 320 °C with a sheath gas rate of 35, aux gas of 10, and max spray current of 100 μA. The full MS scan for both polarities utilized 120,000 resolution with an AGC target of 3e6 and a maximum IT of 100 ms, and the scan range was from 67-1000 m/z. Tandem MS spectra for both positive and negative modes used a resolution of 15,000, AGC target of 1e5, maximum IT of 50 ms, isolation window of 0.4 m/z, isolation offset of 0.1 m/z, fixed first mass of 50 m/z, and 3-way multiplexed normalized collision energies (nCE) of 10, 35, 80. The minimum AGC target was 1e4 with an intensity threshold of 2e5. All data were acquired in profile mode.

*Relative quantification of metabolites.* The resulting Thermo^TM^ RAW files were converted to mzXML format using ReAdW.exe version 4.3.1 to enable peak detection and quantification. The centroided data were searched using an in-house python script Mighty_skeleton version 0.0.2 and peak heights were extracted from the mzXML files based on a previously established library of metabolite retention times and accurate masses adapted from the Whitehead Institute (39), and verified with high-resolution MS/MS spectral curated against the NIST14MS/MS (40) and METLIN (2017) (41) tandem mass spectral libraries. Metabolite peaks were extracted based on the theoretical *m*/*z* of the expected ion type e.g., [M+H]^+^, with a ±5 part-per-million (ppm) tolerance, and a ± 7.5 second peak apex retention time tolerance within an initial retention time search window of ± 0.5 min across the study samples. The resulting data matrix of metabolite intensities for all samples and blank controls was processed with an in-house statistical pipeline Metabolyze version 1.0 and final peak detection was calculated based on a signal to noise ratio (S/N) of 3X compared to blank controls, with a floor of 10,000 (arbitrary units). For samples where the peak intensity was lower than the blank threshold, metabolites were annotated as not detected, and the threshold value was imputed for any statistical comparisons to enable an estimate of the fold change as applicable. The resulting blank corrected data matrix was then used for all group-wise comparisons, and t-tests were performed with the Python SciPy (1.1.0) (42) library to test for differences and generate statistics for downstream analyses. Any metabolite with a p-value < 0.05 was considered significantly regulated (up or down). Heatmaps were generated with hierarchical clustering performed on the imputed matrix values utilizing the R library pheatmap (1.0.12) (43).

***Metagenomic analysis***

Publicly available shotgun metagenomic sequencing data of 395 samples from an observational cohort of hematopoietic stem cell transplant patients (44, 45) was downloaded using the SRA toolkit (46) v. 2.10.9. Kraken (47) v. 2.1.3 with the standard database from October 10, 2023 and default parameters was used for taxonomic classification of reads. Subsequently, the subset of *Staphylococcus* reads was created for each sample using extract_kraken_reads.py with taxonomic ID 1279 and the --include-children flag. These subsets were assembled using spades (48) v. 3.15.5 and the --meta flag; subsequently re-classified with kraken, and filtered to extract *Staphylococcus aureus* contigs with extract_kraken_reads.py using taxonomic ID 1280 and the --include-children flag. From the 395 downloaded metagenomic read sets, 374 produced between ~200 bp and ~68 kbp and one (ID: FMT.0034I) produced 1.1 Mbp of assembled sequence classified as *S. aureus*. A nucleotide collection of the coding sequences of the genes of interest was extracted from NCBI annotated assembly GCA_000013465.1 (genbank format) using BioPython (49) v. 1.78. A second nucleotide collection was constructed with the putative promoter regions of the genes of interest, consisting of the intergenic region upstream of each gene (up to a maximum of 500 bp), additionally including 500 bp of the gene sequence to improve specificity in sequence similarity search. Each of these two sequence collections were used as a query in a blastn (31) v 2.11.0 search, with the *S. aureus* assembled sequence from FMT.0034I as the database, filtering for a minimum identity of 90% and minimum query coverage per high-scoring pair of 70%. The sequence of all blast hits was extracted from FMT.0034I and aligned to their references using pymummer (50) v. 0.11.0 using a minimum identity of 90%, a minimum length of 20, and a breaklength of 200 and the max_match, show_snps, and show_snps_C flags. As indicated in the text, we detected non-synonymous variant alleles in one sample whose metagenomic assemblies included more than 1 Mbp of *S. aureus* nucleotide sequence, allowing for the assembly of candidate genes albeit at a shallow depth of coverage. Matching mutations support the reliability of our sequencing results despite their shallow depth. Nucleotide variants from the pymummer alignment were extracted for all hits, while those in coding sequences were additionally translated into amino-acid changes, considering substitutions, deletions, truncations, and frame-shifts.

***Statistical analysis***

Prism 9 software (GraphPad, Inc.) was used for statistical analyses unless otherwise noted. Precision measures, n values, and statistical tests used are indicated in all figure legends. Symbols: ns = not significant, **P* < 0.05, ** *P* < 0.01, *** *P* < 0.001, **** *P* < 0.0001.

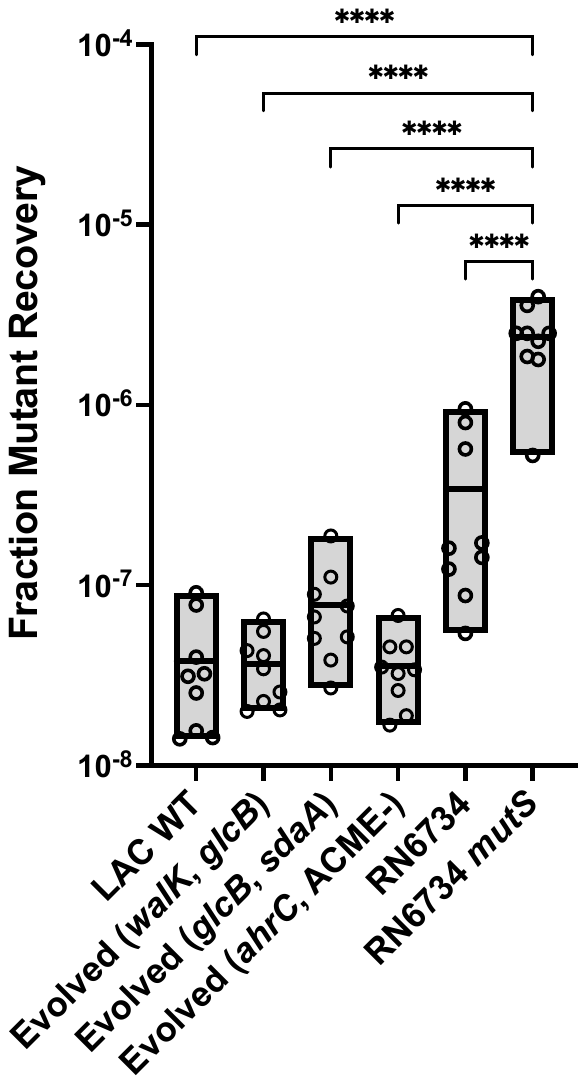

**Fig. S1. Evolved mutants are not hypermutable.** (Related to Fig. 2). Mutation frequency to rifampicin resistance. Evolved strains *walK*, *glcB* (W5-9), *glcB*, *sdaA* (W5-10), *ahrC*, ACME (-) (W5-1-O18), strain LAC (BS819), or control strains RN6734 wild-type and hypermutable *mutS* RN6734 mutant(22) were applied to agar plates containing 40 ng/mL rifampin. Fraction mutant recovery = no. of rifampin-resistant mutants / total no. of colonies. Heavy lines in boxes indicate the mean values. One-way ANOVA, Sidak’s multiple comparisons test: ns *P* > 0.05, **** *P* < 0.0001.

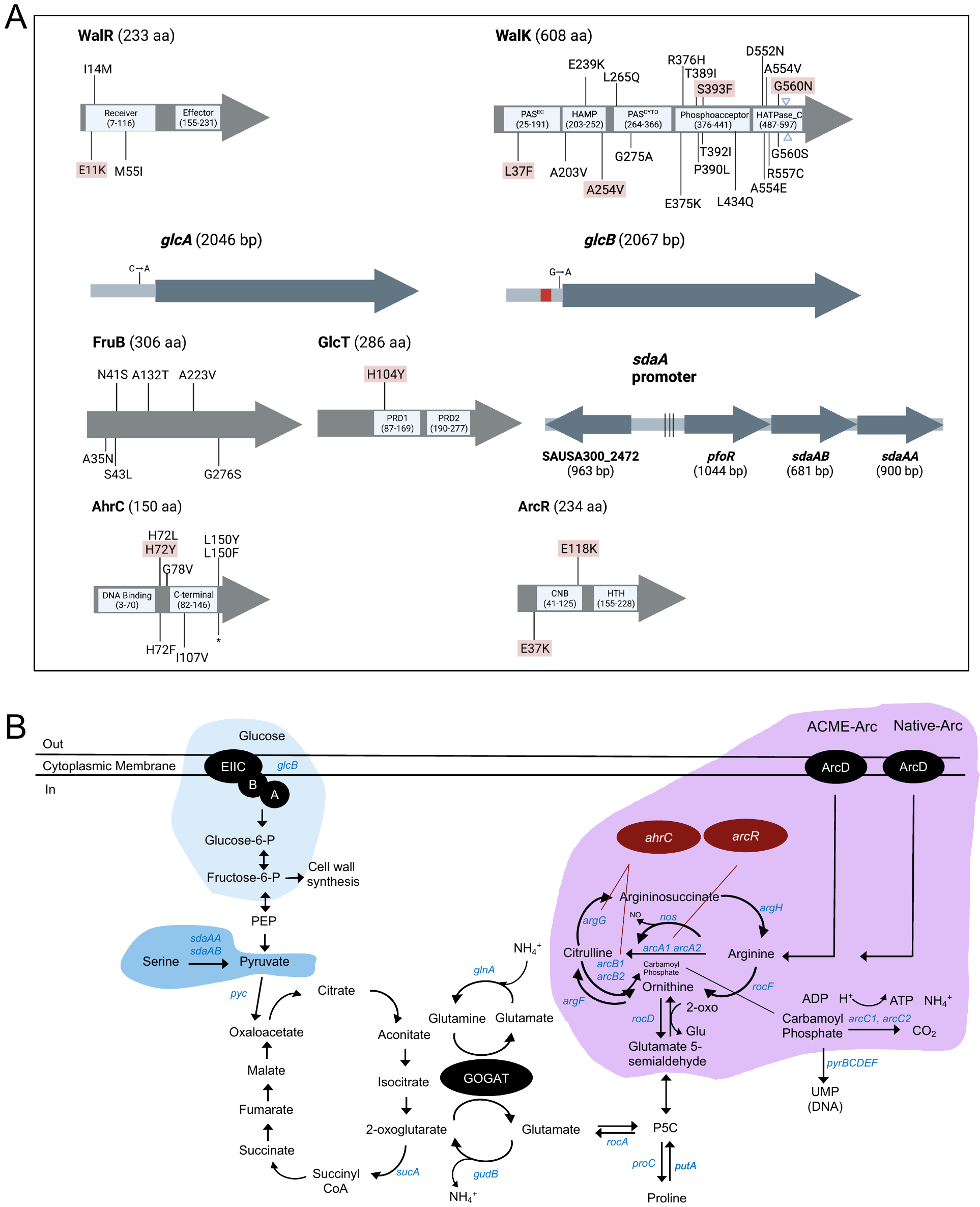

**Fig. S2.** **Mutations in selected genes identified during intestinal adaptation of *S. aureus* in germ-free mice**. (Related to Fig.2). (A) Locations of evolved variation in the indicated genes. Shown are positions of polymorphic nucleotides, amino acid substitutions, insertions (triangles), deletions (red block), and stop-codon mutations (star). Pink color indicates single-nucleotide variants present in isolates from multiple cages. Protein domains were defined as in(13) for WalR and WalK), and the KEGG database (<https://www.genome.jp>) for GlcT, AhrC, and ArcR. (B) Metabolism in *S. aureus* affected by mutant genes Blue color indicates intragenic or intergenic mutations found in evolved isolates. (ACME: arginine catabolic mobile element. P5C: Δ^1^-Pyrroline-5-carboxylate. GOGAT: glutamine oxoglutarate aminotransferase. PEP: phosphoenolpyruvate). Adapted from(51).

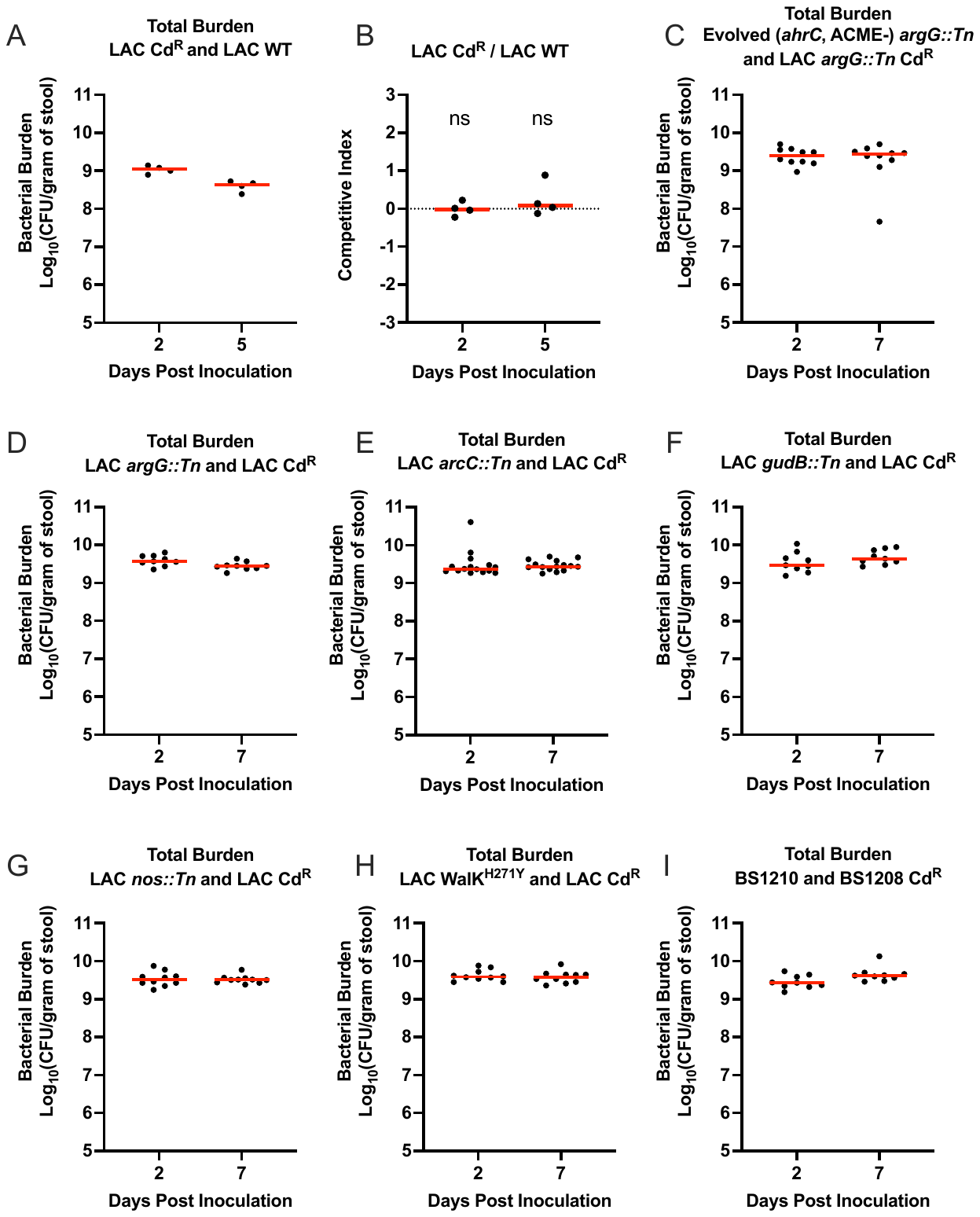

**Fig. S3. Bacterial burden and cadmium resistance marker for competition assays of *S. aureus*** **colonization fitness.** (Related to Figs. 3, 4, and 6). (A-B) Cadmium resistance marker does not affect *S. aureus* colonization fitness in the gut of germ-free mice. Competition assays in germ-free mice involving strain LAC (BS819) and LAC Cd^R^ (VJT32.58). LAC contained a chromosomally integrated cadmium resistance marker (SaPI1 *att*C::*cadCA;* strain VJT32.58) to distinguish the strains following plating of serial dilutions on tryptic soy agar (TSA) with or without cadmium (0.3 mM). Evolved mutants and LAC Cd^R^ were mixed 1:1 and used to inoculate each mouse. Each symbol represents data from one mouse (*n* = 6-9 mice). The red lines are medians. Each symbol represents data from one mouse (*n* = 4 mice). ns *P* > 0.05 by Wilcoxon signed-rank test. (C-I) Bacterial burden results for *in vivo* competition experiments shown in Figs. 4 and 6 (*n* = 9-15 mice). The red lines are medians.

**Fig. S4. Evolved mutations in *glcB* and *glcT* are predicted to increase glucose import.** (Related to Fig. 4). (A) Schematic representation of evolved point mutations upstream of the *glcB* start site (ATG; marked with a rectangular box) and alignment of the *glcB* primary sequences from LAC wild-type (WT) and various evolved mutants. Single nucleotide change (G to A) is highlighted in red. (B) Proposed regulatory mechanism of *glcB* transcriptional activity by GlcT in WT and the evolved mutant GlcT^H104Y^. In the presence of glucose, or a glcT mutation replacing the presumptive regulatory histidine-104, dimeric GlcT stabilizes the antiterminator (active GlcT) in the 5′ UTR of *glcB* mRNA, thereby enabling transcription of the full-length *ptsG* mRNA. In the absence of glucose, PtsG phosphorylates GlcT, preventing it from forming dimers and from RNA binding. Consequently, the 5′ UTR folds into the more stable transcription terminator configuration, causing premature termination of operon transcription. (C) Mfold predictions (37) of the effect of various evolved *glcB* mutations in (A) on the terminator structure of *glcB*. Minimum free energy predictions (dG) indicate that *glcB* terminator loop structures of variants are less stable than those of WT. Location of variant deletions are marked in the WT structure. The putative RAT (ribonucleic antiterminator) sequence is highlighted in blue.

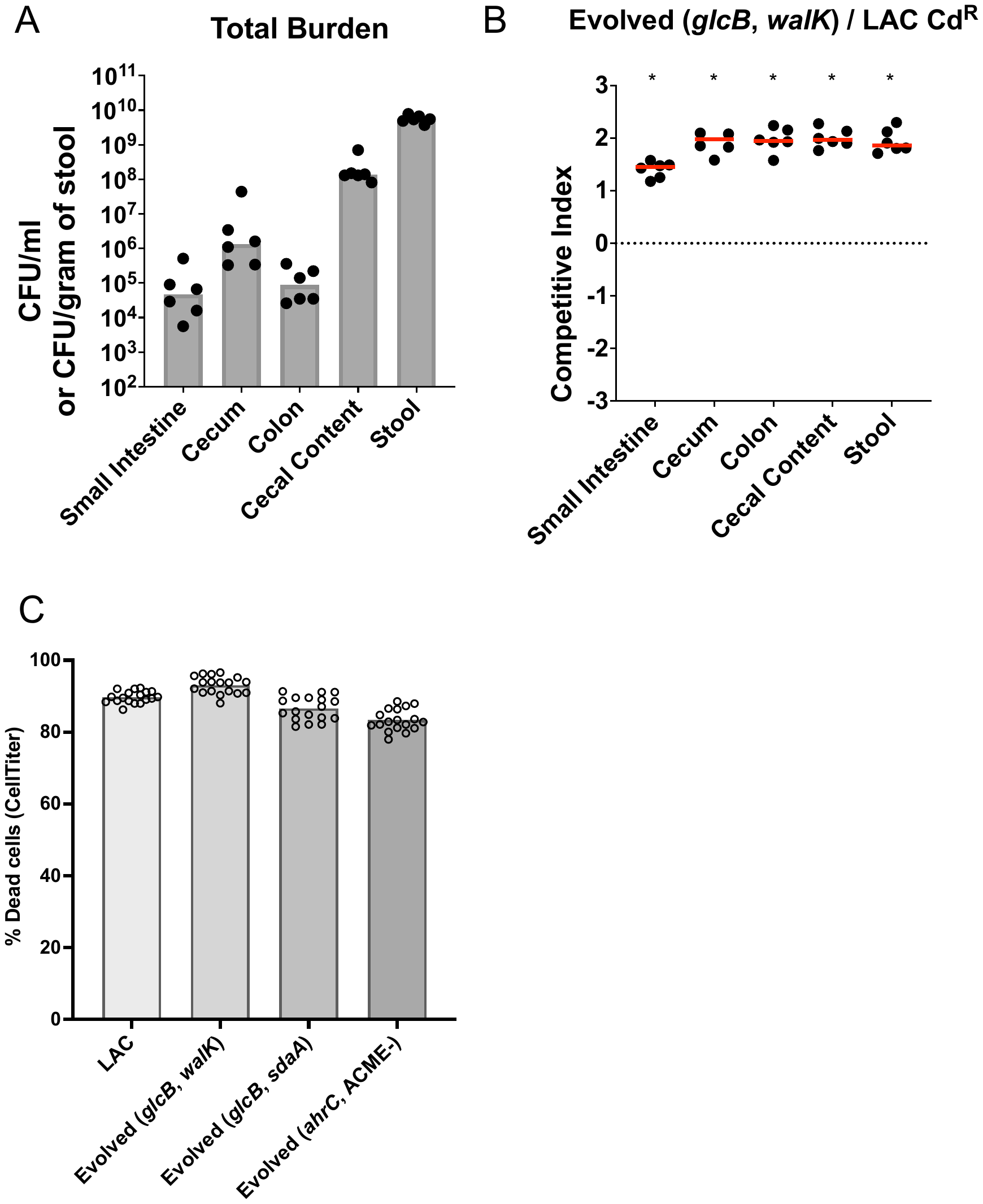

**Fig. S5. *glcB* *walK* mutant demonstrates increased fitness for colonization throughout the gut.** (Related to Figs. 6 and 7). (A-B) Competition assay in germ-free mice, performed as in Fig. S3, involving strains evolved mutant *walK*, evolved mutant *glcB* (W5-9), and control strain LAC Cd^R^ (VJT32.58). Bacteria were quantified from stool and from the indicated GI tract tissues, harvested 6 days after inoculation. Each symbol represents data from one mouse (n = 6 mice). * *P* < 0.05 by Wilcoxon signed-rank test. The red lines are medians. **Evolved changes have limited impact on cytotoxicity compared to parental strain LAC**. (C) Toxicity with primary human neutrophils using 10% (by final volume) filtered culture supernatants. Overnight cultures were diluted and grown for 5 h. Strains, which were used in mouse skin abscess model experiments (Fig. 7), were evolved mutants *glcB*, *walK* (W5-9), *glcB*, *sdaA* (W5-10), and *ahrC*, ACME (-) (W5-1-O18). Each isolate was tested with 2 biological replicates from 3 donors over 2 independent experiments. Cell viability of hPMN cells was measured with the metabolic dye CellTiter. Bars indicate means.

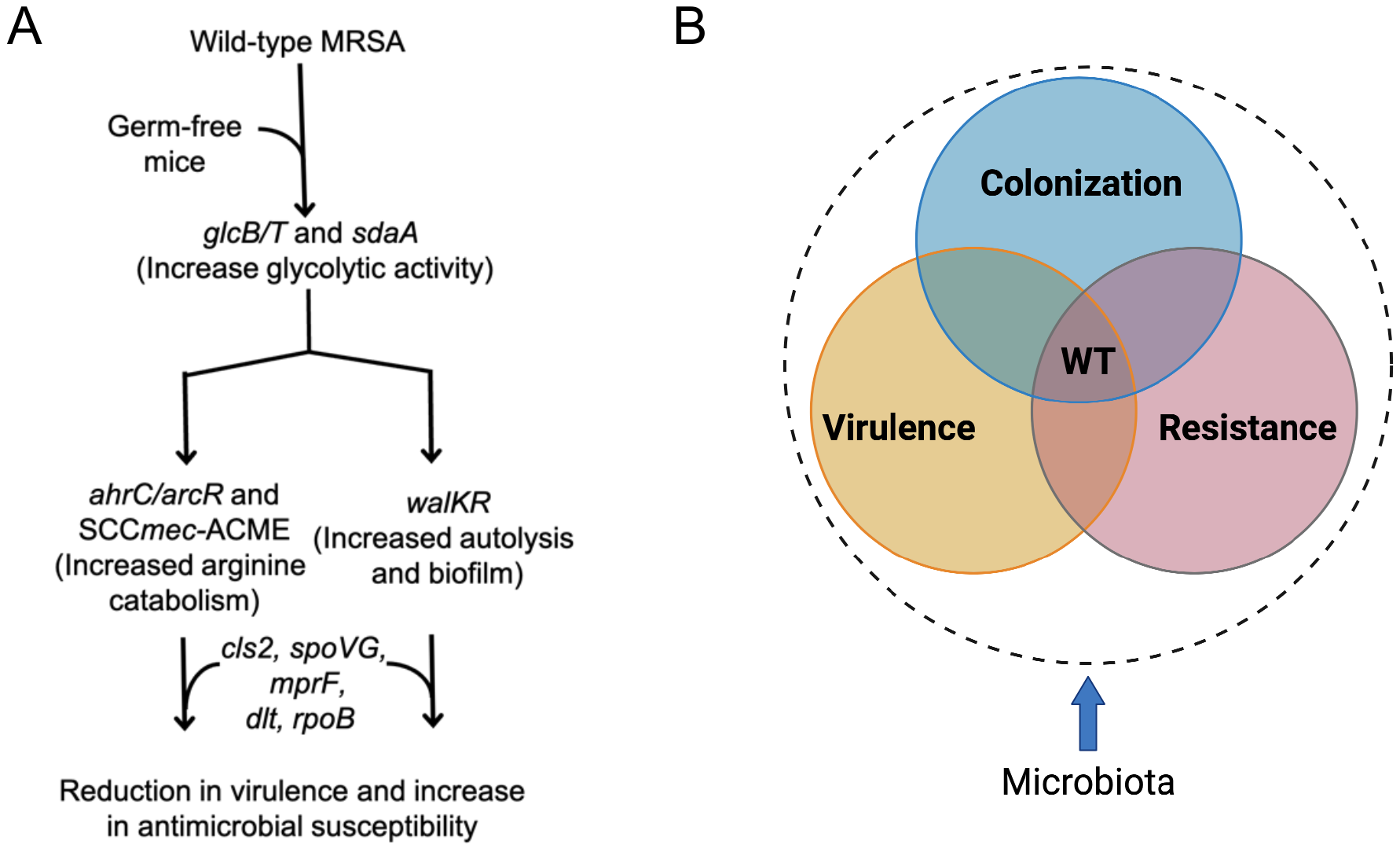

**Fig. S6.** (A) Major steps in metabolic adaptation during intestinal colonization of germ-free mice. Genomic analyses revealed the general rule that primary differentiation of *glcB/glcT* is followed by independent evolution of polymorphism of either *ahrC/arcR* or *walKR*. Predominant members of these two mutational tracks generate either growth or biofilm agonist activities that are potentially driven by intraclonal competition. Antibiotic resistance loci (e.g., *cls2*, *spoVG*) are compromised during colonization adaptation. L-serine catabolism involving *sdaA* is a potential therapeutic target in Enterobacteriaceae to prevent blooming in inflamed intestine (52). Mutations in *ahrC* increase arginine catabolism, especially via *nos*, which has been described previously as an important nasal colonization factor in mice (53). Likewise, *walKR* is specifically expressed in *S. aureus* during nasal colonization in humans (54). The essential nature of the *walKR* system makes it an attractive target for novel antimicrobial compounds that could inhibit *S. aureus* growth (55). Collectively, our results extend the influence of arginine catabolism and *walKR* from nasal to intestinal colonization; they also provide support for three potentially targetable bacterial factors driving growth and colonization fitness. (B) Fitness of adaptive variants as the outcome of a dynamic interaction among colonization, virulence, and resistance, depective through a Venn diagram. The gradients of the three traits interact in complex ways, influencing the fitness outcome at the diagram's center, where they intersect. In this model, the presence of commensal microbiota shapes and constrains MRSA adaptation. Optimal long-term fitness is achieved where colonization, virulence, and resistance traits converge on a wild-type phenotype that is maintained by the microbiota. Critically, fitness outcomes can vary dramatically in different environments, especially if key traits or the host-microbiota relationship changes. Although mutations may provide short-term advantages to MRSA during infection and antibiotic treatment, commensal competitors tend to disadvantage these mutants over the long term during colonization.

**Table S1**. **List of LAC evolved strains used in cefoxitin and daptomycin Etests. Related to Fig. 7C-7E.**

| Strain | Description | Source |
| --- | --- | --- |
| W5-1-O10 | *mec*(-) strain used in cefoxitin Etest | This study |
| W5-1-O11 | *mec*(-) strain used in cefoxitin Etest | This study |
| W5-1-O12 | *mec*(-) strain used in cefoxitin Etest | This study |
| W5-2-O12 | *mec*(-) strain used in cefoxitin Etest | This study |
| W5-3-O3 | *mec*(-) strain used in cefoxitin Etest | This study |
| W5-3-O14 | *mec*(-) strain used in cefoxitin Etest | This study |
| W5-4-O11 | *mec*(-) strain used in cefoxitin Etest | This study |
| W5-1-T13 | *mec*(-) strain used in cefoxitin Etest | This study |
| W5-1-T14 | *mec*(-) strain used in cefoxitin Etest | This study |
| W5-9 | *mec*(-) strain used in cefoxitin Etest | This study |
| W5-10 | *mec*(-) strain used in cefoxitin Etest | This study |
| W5-1-O4 | *mec*(+) strain used in cefoxitin Etest | This study |
| W5-1-O17 | *mec*(+) strain used in cefoxitin Etest | This study |
| W5-2-O3 | *mec*(+) strain used in cefoxitin Etest | This study |
| W5-2-T15 | *mec*(+) strain used in cefoxitin Etest | This study |
| W5-3-T19 | *mec*(+) strain used in cefoxitin Etest | This study |
| W5-4-O12 | *mec*(+) strain used in cefoxitin Etest | This study |
| W5-4-T1 | *mec*(+) strain used in cefoxitin Etest | This study |
| W5-25 | *mec*(+) strain used in cefoxitin Etest | This study |
| W5-1-O18 | *mec*(+) strain used in cefoxitin Etest | This study |
| W5-3-O6 | *mec*(+) strain used in cefoxitin Etest | This study |
| W5-30 | *spoVG* mutant strain used in cefoxitin Etest | This study |
| W5-3-O1 | *spoVG* mutant strain used in cefoxitin Etest | This study |
| W5-3-O8 | *spoVG* mutant strain used in cefoxitin Etest | This study |
| W5-3-O11 | *spoVG* mutant strain used in cefoxitin Etest | This study |
| W5-3-O17 | *spoVG* mutant strain used in cefoxitin Etest | This study |
| W5-25 | *spoVG* mutant strain used in cefoxitin Etest | This study |
| W5-26 | *spoVG* mutant strain used in cefoxitin Etest | This study |
| W5-3-T6 | *spoVG* mutant strain used in cefoxitin Etest | This study |
| W5-3-T7 | *spoVG* mutant strain used in cefoxitin Etest | This study |
| W5-3-T15 | *spoVG* mutant strain used in cefoxitin Etest | This study |
| W5-1-O4 | Non-*spoVG* mutant strain used in cefoxitin Etest | This study |
| W5-1-O13 | Non-*spoVG* mutant strain used in cefoxitin Etest | This study |
| W5-2-O18 | Non-*spoVG* mutant strain used in cefoxitin Etest | This study |
| W5-2-O19 | Non-*spoVG* mutant strain used in cefoxitin Etest | This study |
| W5-3-O4 | Non-*spoVG* mutant strain used in cefoxitin Etest | This study |
| W5-3-O5 | Non-*spoVG* mutant strain used in cefoxitin Etest | This study |
| W5-1-T2 | *cls2* mutant strain used in daptomycin Etest | This study |
| W5-1-T18 | *cls2* mutant strain used in daptomycin Etest | This study |
| W5-1-T19 | *cls2* mutant strain used in daptomycin Etest | This study |
| W5-1-T20 | *cls2* mutant strain used in daptomycin Etest | This study |
| W5-2-T2 | *cls2* mutant strain used in daptomycin Etest | This study |
| W5-2-T4 | *cls2* mutant strain used in daptomycin Etest | This study |
| W5-2-T10 | *cls2* mutant strain used in daptomycin Etest | This study |
| W5-2-T18 | *cls2* mutant strain used in daptomycin Etest | This study |
| W5-1-T16 | *cls2* mutant strain used in daptomycin Etest | This study |
| W5-1-T9 | *cls2* mutant strain used in daptomycin Etest | This study |
| W5-1-T10 | *cls2* mutant strain used in daptomycin Etest | This study |
| W5-1-T15 | *cls2* mutant strain used in daptomycin Etest | This study |
| W5-5 | *cls2* mutant strain used in daptomycin Etest | This study |
| W5-8 | *cls2* mutant strain used in daptomycin Etest | This study |
| W5-9 | *cls2* mutant strain used in daptomycin Etest | This study |
| W5-1-O15 | Non*-cls2* mutant strain used in daptomycin Etest | This study |
| W5-1-O17 | Non*-cls2* mutant strain used in daptomycin Etest | This study |
| W5-2-O1 | Non*-cls2* mutant strain used in daptomycin Etest | This study |
| W5-4-O6 | Non*-cls2* mutant strain used in daptomycin Etest | This study |
| W5-4-O16 | Non*-cls2* mutant strain used in daptomycin Etest | This study |
| W5-3-O17 | Non*-cls2* mutant strain used in daptomycin Etest | This study |
| W5-3-O16 | Non*-cls2* mutant strain used in daptomycin Etest | This study |

**Table S2. List of bacterial strains used in this study.**

| Strain | Source |
| --- | --- |
| LAC (*S. aureus* USA300 wild-type; AH1263; BS819) | Boles et al (56). |
| LAC Cd^R^ (VJT 32.58) | Dumont et al (12). |
| LAC *Δatl::tet* (VJT 80.33) | Zheng et al (57). |
| LAC *argG::bursa* (BS1313) | This study |
| LAC *argG::bursa* Cd^R^ (BS1539) | This study |
| LAC *arcC::bursa* (BS1535) | This study |
| LAC *gudB::bursa* (BS1409) | This study |
| LAC *nos:: bursa* (BS1434) | This study |
| LAC *mprF::bursa* (VJT57.49; BS1328) | This study |
| LAC WalK^H271Y^ (BS1274) | This study |
| BS1565 (LAC evolved week 1 isolate) | This study |
| LAC Evolved W3-1C (week 3 isolate) | This study |
| LAC Evolved W5-9 | This study |
| LAC Evolved W5-10 | This study |
| LAC Evolved W5-1-O18 | This study |
| LAC Evolved W5-17 | This study |
| LAC Evolved W5-1-O1 | This study |
| LAC Evolved W5-7 | This study |
| LAC Evolved W5-2-O10 | This study |
| LAC Evolved W5-1-O18 *argG:: bursa* (BS1541) | This study |
| LAC Evolved W5-25 | This study |
| LAC Evolved W5-2 | This study |
| LAC Evolved W5-21 | This study |
| LAC Evolved W5-2-T2 | This study |
| LAC Evolved W5-2-T6 | This study |
| LAC Evolved W5-4-T16 | This study |
| For evolved strains used in cefoxitin and daptomycin Etests, see Table S1. | This study |
| JE2 *argG::bursa* | Fey et al (11) |
| JE2 *arcC::bursa* | Fey et al (11). |
| JE2 *gudB::bursa* | Fey et al (11). |
| JE2 *nos::bursa* | Fey et al (11). |
| JE2 *mecA::bursa* (BS1168) | Fey et al (11). |
| JH1 | Mwangi et al (58). |
| RN6734 | Altman et al (59). |
| RN6734 *mutS* | Altman et al (59). |
| BS1208 (VSSA clinical isolate JH1) | Mwangi et al (23). |
| BS1208 Cd^R^ (BS1709) | This study |
| BS1210 (hVISA clinical isolate, *yycH* mutant) | This study |
| Mu3 (hVISA, BS626) | Ohta et al (60). |
| *Escherichia coli* IM08B | Monk et al (1). |

**Table S3. List of oligonucleotides used in this study.**

| Oligonucleotide | Source |
| --- | --- |
| E239K_scr_F: GTCATCCTAGGATTCTTTATAGCG | This paper |
| walRK_F2:  ATTTCCTCCAACAACATGAG | This paper |
| walRK_R1:  TTATTATTCATCCCAATCACC | This paper |
| ptsG_F3:  CTATGCACCAGGTATCGGTCAAATC | This paper |
| ptsG_R3:  CACTGTACCATCAAACGGTGCTAC | This paper |
| 16S_rRNA_F: ACGTGGATAACCTACCTATAAGACTGGGAT | Delaune et al (16). |
| 16S_rRNA_R: TACCTTACCAACTAGCTAATGCAGCG | Delaune et al (16). |

**References**

1.	Monk IR, Tree JJ, Howden BP, Stinear TP, Foster TJ. 2015. Complete bypass of restriction systems for major *Staphylococcus aureus* lineages. mBio 6:e00308-15.

2.	Hussain M, Hastings JG, White PJ. 1991. A chemically defined medium for slime production by coagulase-negative staphylococci. J Med Microbiol 34:143-7.

3.	Hussain M, Hastings JGM, White PJ. 1991. A chemically defined medium for slime production by coagulase-negative staphylococci. Journal of Medical Microbiology 34:143-147.

4.	Marchiando AM, Ramanan D, Ding Y, Gomez LE, Hubbard-Lucey VM, Maurer K, Wang C, Ziel JW, van Rooijen N, Nunez G, Finlay BB, Mysorekar IU, Cadwell K. 2013. A deficiency in the autophagy gene Atg16L1 enhances resistance to enteric bacterial infection. Cell Host Microbe 14:216-24.

5.	Brugiroux S, Beutler M, Pfann C, Garzetti D, Ruscheweyh HJ, Ring D, Diehl M, Herp S, Lotscher Y, Hussain S, Bunk B, Pukall R, Huson DH, Munch PC, McHardy AC, McCoy KD, Macpherson AJ, Loy A, Clavel T, Berry D, Stecher B. 2016. Genome-guided design of a defined mouse microbiota that confers colonization resistance against *Salmonella enterica* serovar Typhimurium. Nat Microbiol 2:16215.

6.	Dallari S, Heaney T, Rosas-Villegas A, Neil JA, Wong SY, Brown JJ, Urbanek K, Herrmann C, Depledge DP, Dermody TS, Cadwell K. 2021. Enteric viruses evoke broad host immune responses resembling those elicited by the bacterial microbiome. Cell Host Microbe 29:1014-1029 e8.

7.	Gries DM, Pultz NJ, Donskey CJ. 2005. Growth in cecal mucus facilitates colonization of the mouse intestinal tract by methicillin-resistant *Staphylococcus aureus*. J Infect Dis 192:1621-7.

8.	Copin R, Sause WE, Fulmer Y, Balasubramanian D, Dyzenhaus S, Ahmed JM, Kumar K, Lees J, Stachel A, Fisher JC, Drlica K, Phillips M, Weiser JN, Planet PJ, Uhlemann AC, Altman DR, Sebra R, van Bakel H, Lighter J, Torres VJ, Shopsin B. 2019. Sequential evolution of virulence and resistance during clonal spread of community-acquired methicillin-resistant *Staphylococcus aureus*. Proc Natl Acad Sci U S A doi:10.1073/pnas.1814265116.

9.	Schneider CA, Rasband WS, Eliceiri KW. 2012. NIH Image to ImageJ: 25 years of image analysis. Nat Methods 9:671-5.

10.	Leech JM, Lacey KA, Mulcahy ME, Medina E, McLoughlin RM. 2017. IL-10 plays opposing roles during *Staphylococcus aureus* systemic and localized infections. J Immunol doi:10.4049/jimmunol.1601018.

11.	Fey PD, Endres JL, Yajjala VK, Widhelm TJ, Boissy RJ, Bose JL, Bayles KW. 2013. A genetic resource for rapid and comprehensive phenotype screening of nonessential *Staphylococcus aureus* genes. MBio 4:e00537-12.

12.	DuMont AL, Yoong P, Surewaard BG, Benson MA, Nijland R, van Strijp JA, Torres VJ. 2013. *Staphylococcus aureus* elaborates leukocidin AB to mediate escape from within human neutrophils. Infect Immun 81:1830-41.

13.	Monk IR, Shaikh N, Begg SL, Gajdiss M, Sharkey LKR, Lee JYH, Pidot SJ, Seemann T, Kuiper M, Winnen B, Hvorup R, Collins BM, Bierbaum G, Udagedara SR, Morey JR, Pulyani N, Howden BP, Maher MJ, McDevitt CA, King GF, Stinear TP. 2019. Zinc-binding to the cytoplasmic PAS domain regulates the essential WalK histidine kinase of *Staphylococcus aureus*. Nature Communications 10:3067.

14.	Schuster CF, Howard SA, Gründling A. 2019. Use of the counter selectable marker PheS* for genome engineering in *Staphylococcus aureus*. Microbiology (Reading) 165:572-584.

15.	Monk IR, Shah IM, Xu M, Tan M-W, Foster TJ. 2012. Transforming the untransformable: application of direct transformation to manipulate genetically *Staphylococcus aureus* and *Staphylococcus epidermidis*. mBio 3:e00277-11.

16.	Delauné A, Dubrac S, Blanchet C, Poupel O, Mäder U, Hiron A, Leduc A, Fitting C, Nicolas P, Cavaillon J-M, Adib-Conquy M, Msadek T. 2012. The WalKR system controls major staphylococcal virulence genes and is involved in triggering the host inflammatory response. Infection and Immunity 80:3438-3453.

17.	Livak KJ, Schmittgen TD. 2001. Analysis of relative gene expression data using real-time quantitative PCR and the 2(-Delta Delta C(T)) Method. Methods 25:402-8.

18.	Lacey KA, Serpas L, Makita S, Wang Y, Rashidfarrokhi A, Soni C, Gonzalez S, Moreira A, Torres VJ, Reizis B. 2023. Secreted mammalian DNases protect against systemic bacterial infection by digesting biofilms. J Exp Med 220.

19.	Yusof A, Engelhardt A, Karlsson A, Bylund L, Vidh P, Mills K, Wootton M, Walsh TR. 2008. Evaluation of a new Etest vancomycin-teicoplanin strip for detection of glycopeptide-intermediate *Staphylococcus aureus* (GISA), in particular, heterogeneous GISA. J Clin Microbiol 46:3042-7.

20.	Hafer C, Lin Y, Kornblum J, Lowy FD, Uhlemann AC. 2012. Contribution of selected gene mutations to resistance in clinical isolates of vancomycin-intermediate *Staphylococcus aureus*. Antimicrob Agents Chemother 56:5845-51.

21.	Hiramatsu K, Aritaka N, Hanaki H, Kawasaki S, Hosoda Y, Hori S, Fukuchi Y, Kobayashi I. 1997. Dissemination in Japanese hospitals of strains of *Staphylococcus aureus* heterogeneously resistant to vancomycin. Lancet 350:1670-3.

22.	O'Neill AJ, Chopra I. 2002. Insertional inactivation of mutS in *Staphylococcus aureus* reveals potential for elevated mutation frequencies, although the prevalence of mutators in clinical isolates is low. J Antimicrob Chemother 50:161-9.

23.	Balasubramanian D, Ohneck EA, Chapman J, Weiss A, Kim MK, Reyes-Robles T, Zhong J, Shaw LN, Lun DS, Ueberheide B, Shopsin B, Torres VJ. 2016. *Staphylococcus aureus* coordinates leukocidin expression and pathogenesis by sensing metabolic fluxes via RpiRc. mBio 7.

24.	Walker BJ, Abeel T, Shea T, Priest M, Abouelliel A, Sakthikumar S, Cuomo CA, Zeng Q, Wortman J, Young SK, Earl AM. 2014. Pilon: An integrated tool for comprehensive microbial variant detection and genome assembly improvement. PLOS ONE 9:e112963.

25.	Li W, O'Neill KR, Haft DH, DiCuccio M, Chetvernin V, Badretdin A, Coulouris G, Chitsaz F, Derbyshire MK, Durkin AS, Gonzales NR, Gwadz M, Lanczycki CJ, Song JS, Thanki N, Wang J, Yamashita RA, Yang M, Zheng C, Marchler-Bauer A, Thibaud-Nissen F. 2021. RefSeq: expanding the Prokaryotic Genome Annotation Pipeline reach with protein family model curation. Nucleic Acids Res 49:D1020-d1028.

26.	Ponstingl H, Ning Z. 2010. SMALT - A new mapper for DNA sequencing reads, vol 1.

27.	Danecek P, Bonfield JK, Liddle J, Marshall J, Ohan V, Pollard MO, Whitwham A, Keane T, McCarthy SA, Davies RM, Li H. 2021. Twelve years of SAMtools and BCFtools. Gigascience 10.

28.	Bush SJ, Foster D, Eyre DW, Clark EL, De Maio N, Shaw LP, Stoesser N, Peto TEA, Crook DW, Walker AS. 2020. Genomic diversity affects the accuracy of bacterial single-nucleotide polymorphism–calling pipelines. GigaScience 9.

29.	Garrison E, Marth G. 2012. Haplotype-based variant detection from short-read sequencing. arXiv preprint arXiv:12073907.

30.	Cingolani P, Platts A, Wang le L, Coon M, Nguyen T, Wang L, Land SJ, Lu X, Ruden DM. 2012. A program for annotating and predicting the effects of single nucleotide polymorphisms, SnpEff: SNPs in the genome of *Drosophila melanogaster* strain w1118; iso-2; iso-3. Fly (Austin) 6:80-92.

31.	Altschul SF, Gish W, Miller W, Myers EW, Lipman DJ. 1990. Basic local alignment search tool. Journal of Molecular Biology 215:403-410.

32.	Hunt M, Mather AE, Sanchez-Buso L, Page AJ, Parkhill J, Keane JA, Harris SR. 2017. ARIBA: rapid antimicrobial resistance genotyping directly from sequencing reads. Microb Genom 3:e000131.

33.	Stamatakis A. 2014. RAxML version 8: a tool for phylogenetic analysis and post-analysis of large phylogenies. Bioinformatics 30:1312-1313.

34.	R Development Core Team. 2010. R: A language and environment for statistical computing, R Foundation for Statistical Computing, <https://www.R-project.org/>.

35.	Letunic I, Bork P. 2021. Interactive Tree Of Life (iTOL) v5: an online tool for phylogenetic tree display and annotation. Nucleic Acids Research 49:W293-W296.

36.	Langbein I, Bachem S, Stulke J. 1999. Specific interaction of the RNA-binding domain of the *Bacillus subtilis* transcriptional antiterminator GlcT with its RNA target, RAT. J Mol Biol 293:795-805.

37.	Zuker M. 2003. Mfold web server for nucleic acid folding and hybridization prediction. Nucleic Acids Research 31:3406-3415.

38.	Jones DR, Wu Z, Chauhan D, Anderson KC, Peng J. 2014. A nano ultra-performance liquid chromatography-high resolution mass spectrometry approach for global metabolomic profiling and case study on drug-resistant multiple myeloma. Anal Chem 86:3667-75.

39.	Chen WW, Freinkman E, Wang T, Birsoy K, Sabatini DM. 2016. Absolute quantification of matrix metabolites reveals the dynamics of mitochondrial metabolism. Cell 166:1324-1337.e11.

40.	Simón-Manso Y, Lowenthal MS, Kilpatrick LE, Sampson ML, Telu KH, Rudnick PA, Mallard WG, Bearden DW, Schock TB, Tchekhovskoi DV, Blonder N, Yan X, Liang Y, Zheng Y, Wallace WE, Neta P, Phinney KW, Remaley AT, Stein SE. 2013. Metabolite profiling of a NIST Standard Reference Material for human plasma (SRM 1950): GC-MS, LC-MS, NMR, and clinical laboratory analyses, libraries, and web-based resources. Anal Chem 85:11725-31.

41.	Smith CA, O'Maille G, Want EJ, Qin C, Trauger SA, Brandon TR, Custodio DE, Abagyan R, Siuzdak G. 2005. METLIN: a metabolite mass spectral database. Ther Drug Monit 27:747-51.

42.	Virtanen P, Gommers R, Oliphant TE, Haberland M, Reddy T, Cournapeau D, Burovski E, Peterson P, Weckesser W, Bright J, van der Walt SJ, Brett M, Wilson J, Millman KJ, Mayorov N, Nelson ARJ, Jones E, Kern R, Larson E, Carey CJ, Polat İ, Feng Y, Moore EW, VanderPlas J, Laxalde D, Perktold J, Cimrman R, Henriksen I, Quintero EA, Harris CR, Archibald AM, Ribeiro AH, Pedregosa F, van Mulbregt P, Vijaykumar A, Bardelli AP, Rothberg A, Hilboll A, Kloeckner A, Scopatz A, Lee A, Rokem A, Woods CN, Fulton C, Masson C, Häggström C, Fitzgerald C, Nicholson DA, Hagen DR, Pasechnik DV, et al. 2020. SciPy 1.0: fundamental algorithms for scientific computing in Python. Nature Methods 17:261-272.

43.	Kolde R. 2012. Pheatmap: pretty heatmaps. R package version 1:726.

44.	Yan J, Liao C, Taylor BP, Fontana E, Amoretti LA, Wright RJ, Littmann ER, Dai A, Waters N, Peled JU, Taur Y, Perales M-A, Siranosian BA, Bhatt AS, van den Brink MRM, Pamer EG, Schluter J, Xavier JB. 2022. A compilation of fecal microbiome shotgun metagenomics from hematopoietic cell transplantation patients. Scientific Data 9:219.

45.	Liao C. 2020. samples(figshare) doi:<https://doi.org/10.6084/m9.figshare.12016983.v9>.

46.	Leinonen R, Sugawara H, Shumway M. 2011. The sequence read archive. Nucleic Acids Res 39:D19-21.

47.	Wood DE, Lu J, Langmead B. 2019. Improved metagenomic analysis with Kraken 2. Genome Biology 20:257.

48.	Prjibelski A, Antipov D, Meleshko D, Lapidus A, Korobeynikov A. 2020. Using SPAdes De Novo Assembler. Current Protocols in Bioinformatics 70:e102.

49.	Cock PJA, Antao T, Chang JT, Chapman BA, Cox CJ, Dalke A, Friedberg I, Hamelryck T, Kauff F, Wilczynski B, de Hoon MJL. 2009. Biopython: freely available Python tools for computational molecular biology and bioinformatics. Bioinformatics 25:1422-1423.

50.	Hunt M, Mather AE, Sánchez-Busó L, Page AJ, Parkhill J, Keane JA, Harris SR. 2017. ARIBA: rapid antimicrobial resistance genotyping directly from sequencing reads. Microb Genom 3:e000131.

51.	Halsey CR, Lei S, Wax JK, Lehman MK, Nuxoll AS, Steinke L, Sadykov M, Powers R, Fey PD. 2017. Amino acid catabolism in *Staphylococcus aureus* and the function of carbon catabolite repression. mBio 8:e01434-16.

52.	Kitamoto S, Alteri CJ, Rodrigues M, Nagao-Kitamoto H, Sugihara K, Himpsl SD, Bazzi M, Miyoshi M, Nishioka T, Hayashi A, Morhardt TL, Kuffa P, Grasberger H, El-Zaatari M, Bishu S, Ishii C, Hirayama A, Eaton KA, Dogan B, Simpson KW, Inohara N, Mobley HLT, Kao JY, Fukuda S, Barnich N, Kamada N. 2020. Dietary L-serine confers a competitive fitness advantage to *Enterobacteriaceae* in the inflamed gut. Nat Microbiol 5:116-125.

53.	Kinkel TL, Ramos-Montanez S, Pando JM, Tadeo DV, Strom EN, Libby SJ, Fang FC. 2016. An essential role for bacterial nitric oxide synthase in *Staphylococcus aureus* electron transfer and colonization. Nat Microbiol 2:16224.

54.	Burian M, Rautenberg M, Kohler T, Fritz M, Krismer B, Unger C, Hoffmann WH, Peschel A, Wolz C, Goerke C. 2010. Temporal expression of adhesion factors and activity of global regulators during establishment of *Staphylococcus aureus* nasal colonization. J Infect Dis 201:1414-21.

55.	Okada A, Igarashi M, Okajima T, Kinoshita N, Umekita M, Sawa R, Inoue K, Watanabe T, Doi A, Martin A, Quinn J, Nishimura Y, Utsumi R. 2010. Walkmycin B targets WalK (YycG), a histidine kinase essential for bacterial cell growth. J Antibiot (Tokyo) 63:89-94.

56.	Boles BR, Thoendel M, Roth AJ, Horswill AR. 2010. Identification of genes involved in polysaccharide-independent *Staphylococcus aureus* biofilm formation. PLOS ONE 5:e10146.

57.	Zheng X, Ma SX, St John A, Torres VJ. 2022. The major autolysin Atl regulates the virulence of *Staphylococcus aureus* by controlling the sorting of LukAB. Infect Immun 90:e0005622.

58.	Mwangi MM, Wu SW, Zhou Y, Sieradzki K, de Lencastre H, Richardson P, Bruce D, Rubin E, Myers E, Siggia ED, Tomasz A. 2007. Tracking the *in vivo* evolution of multidrug resistance in *Staphylococcus aureus* by whole-genome sequencing. Proceedings of the National Academy of Sciences 104:9451-9456.

59.	Altman DR, Sullivan MJ, Chacko KI, Balasubramanian D, Pak TR, Sause WE, Kumar K, Sebra R, Deikus G, Attie O, Rose H, Lewis M, Fulmer Y, Bashir A, Kasarskis A, Schadt EE, Richardson AR, Torres VJ, Shopsin B, Bakel Hv. 2018. Genome plasticity of *agr*-defective *Staphylococcus aureus* during clinical infection. Infection and Immunity 86:e00331-18.

60.	Ohta T, Hirakawa H, Morikawa K, Maruyama A, Inose Y, Yamashita A, Oshima K, Kuroda M, Hattori M, Hiramatsu K, Kuhara S, Hayashi H. 2004. Nucleotide substitutions in *Staphylococcus aureus* strains, Mu50, Mu3, and N315. DNA Research 11:51-56.
